# Multiscale characterization of *Caenorhabditis elegans* mutants to probe functional mechanisms of human actin pathological variants

**DOI:** 10.1101/2024.07.22.604239

**Authors:** Théo Hecquet, Nadine Arbogast, Delphine Suhner, Anaïs Goetz, Grégory Amann, Selin Yürekli, Fiona Marangoni, Sophie Quintin, Johannes N. Greve, Nataliya Di Donato, Anne-Cécile Reymann

**Affiliations:** Université de Strasbourg, IGBMC UMR 7104-UMR-S 1258, F-67400 Illkirch, France; CNRS, UMR 7104, F-67400 Illkirch, France; Inserm, UMR-S 1258, F-67400 Illkirch, France; IGBMC, Institut de Génétique et de Biologie Moléculaire et Cellulaire, F-67400 Illkirch, France; Institute for Biophysical Chemistry, Hannover Medical School, Fritz–Hartmann–Centre for Medical Research, 30625 Hannover, Germany; Department of Human Genetics, Hannover Medical School, 30§625 Hannover, Germany

**Keywords:** *C. elegans*, cytoplasmic actin, rare disease, Non-Muscle Actinopathy, Baraitser-Winter Cerebrofrontofacial Syndrome

## Abstract

Actin plays a crucial role in diverse physiological processes by forming dynamic networks that determine cellular shape and mechanical properties. Non-Muscle Actinopathies (NMA) are rare diseases caused by *de novo* variants in human cytoskeletal β-actin (*ACTB*) and γ-actin (*ACTG1*) genes, ranging from missense mutations to whole gene deletions. Currently, the high clinical variability and genotype-phenotype correlations in NMA remain largely unresolved. To address this concern, we inserted nine mutations identified in patients in the *C. elegans* cytoplasmic actin orthologue *act-2* and performed a quantitative multiscale characterization of these animal models. We uncovered various perturbations including micro-scale actin network defects, cell-scale abnormalities, morphogenesis failure, and weaker behavioural phenotypes. Notably, the severity of the observed defects correlates with the severity of patients’ symptoms. Thus, we provide evidence that such *C. elegans* models are relevant to investigate the mechanisms underlying NMA physiopathology and could ultimately be used to screen for therapeutic strategies.

## INTRODUCTION

The ability to dynamically assemble the actin cytoskeleton in response to internal and external triggers is a hallmark of a healthy and functional cell. Finely tuned actin architectures are coordinated to fulfil specific cellular needs and drive a multitude of cellular processes including motility, adhesion, morphogenesis, mechanotransduction, internal organisation and trafficking^1–4^. Constant reorganization for adapting to external cues is intrinsically linked to the capacity of cytoskeletal actin networks to turnover and to reorganise, typically in the scale of seconds or minutes^3^. Altered cytoskeletal integrity or dynamics are implicated in many disorders ranging from hearing loss to cancer or to rare developmental diseases^5,6^. The human genome encodes six highly conserved actin isoforms, which are produced in a time- and tissue-specific manner and every actin isoform has been associated with human pathologies^5^. *De novo* missense variants, frameshift or truncating variants as well as gene deletions in the *ACTB* and *ACTG1* genes, which encode the ubiquitously expressed cytoplasmic β- and γ-actin, cause substantial cellular dysfunction and lead to a range of rare autosomal dominant disorders, termed Non-Muscle Actinopathies (NMA)^7–12^. One particularity of this category of diseases is the broad spectrum of symptoms found in patients. NMA can range from very mild to severe neurological impairment and cortical malformations as well as reduced life expectancy of the affected patients. So far, 290 patients and 157 unique mutations have been reported to cause NMA^13^. The most frequently associated NMA disease, named Baraitser-Winter cerebrofrontofacial syndrome (BWCFF), is associated with characteristic facial malformations, intellectual disability, cortical malformations, epilepsy, hearing loss and a variety of congenital anomalies^10,11^. Even within the BWCFF subclass of NMA, a large range of symptom severity is observed.

To date, a definitive genotype-phenotype correlation within the NMA field has yet to emerge, and the functional mechanisms causing symptoms in patients remain largely elusive. It is likely that the complexity originates from both the multifaceted roles of actin and its numerous molecular partners (Actin Binding Proteins; ABP). Furthermore, the existence of compensatory mechanisms between isoforms constitute a challenge in the study of actin *in vivo*^14–19^. Actin variants have been proposed to affect the cytoskeleton by four mechanisms: insufficient availability of actin monomers, alteration of the actin filament assembly or disassembly kinetics, alteration of actin interactions with binding partners and assembly of toxic actin oligomers detrimental for proper cytoskeletal function^13,20^. The use of purified recombinant human actin proteins allows us to quantitatively assess defects in terms of folding, thermal stability, nucleotide-binding, G- to F-actin transition and interaction with ABPs for some variants^13,20–22^, validating some of these hypotheses. At the structural level, *in silico* models correlate some NMA mutations with nucleotide-binding, monomer-monomer interaction and canonical G-actin-ABP interaction surfaces^8^. *In vivo*, the effects of *Actb* and *Actg1* deletions have been studied in mouse embryonic fibroblasts and in the whole animal^23–25^. Notably, *Actb* deletion leads to proliferation defects in mouse embryonic fibroblasts as well as embryonic lethality in mice, whereas *Actg1* null mutant mice are viable^26^. Currently, however, a comprehensive system capable of bridging these different scales and validating the suggested molecular to functional relationship is still missing.

To fill this gap, we chose to use the free-living nematode *Caenorhabditis elegans* (*C. elegans*) as a model to study NMA, as it provides easy access to all biologically relevant scales: from general animal fitness to stereotypical organization and development, up to cell-scale events and single molecular dynamics *in vivo*. *C. elegans* has already been used as a model for numerous diseases, notably for the study of neurodegenerative and neuromuscular disorders^27^, validating its relevance for assessing the molecular-to-functional link in pathologies. The ease of genetic manipulation and the ability to obtain stable, isogenic strains within a matter of weeks streamlines the process of strain characterization and enables us to investigate the consequences of multiple NMA variants in a reasonable amount of time.

As in humans, *C. elegans* relies on different cytoplasmic actin paralogs to sustain cellular cytoskeleton integrity in all cell types. Three ubiquitously expressed actin genes (*act-1, act-2* and *act-3*) are present in a cluster and operate in a partially redundant manner, notably during early embryogenesis^28–30^. These three cytoplasmic *C. elegans* actins are also required for proper muscle function^31^. Indeed, mutants which carry some *act-2* mutations, exhibit uncoordinated adult movement or pharyngeal pumping defects^29,32–34^. A fourth actin-coding gene named *act-4* is expressed in different tissues, notably in neurons and in muscle cells. However, *act-4* is not as abundant or ubiquitously expressed in the early embryo as expected for an essential cytoplasmic actin such as *act-1 -2, and -3*^18^. By contrast, *act-5* shows a strong tissue-specific expression in the gut and excretory cell^35^. Among the *C. elegans* proteins, ACT-2 has the highest similarity to human cytoplasmic actin proteins, sharing 98% of amino acids with β-actin and 97% with γ-actin.

In this study, we assess the consequences of actin variants associated with BWCFF or other NMA in a model organism and present a quantitative multiscale characterization of *C. elegans* mutants. As cytoplasmic actin is required pleiotropically, we performed a global survey of mutants using well-established methods to assay both nematode fitness and specific organ function. We propose a classification of selected NMA variants with respect to the observed phenotypes in *C. elegans* and compare this classification to the severity of human patient’s symptoms. This initial study of actin variants showed a strong correlation in the degree of pathological severities between *C. elegans* and humans, thus validating the use of *C. elegans* as a tool to study NMA. Importantly, this work paves the way towards the prediction of clinical severity in patients carrying novel actin variants from the outset of diagnosis.

## RESULTS

### Generation of C. elegans models recapitulating human non-muscle actinopathies

With respects to the three *C. elegans* cytoplasmic actin genes: while *act-1* and *act-3* share identical coding sequences, they differ from *act-2* by three non-synonymous and many silent variations as well as distinct intronic regions. This allows a slightly easier CRISPR/Cas9 genome editing for *act-2* than for the others^36^. In addition, similarly to its human cytoplasmic actin counterparts (*ACTB* and *ACTG1*), *C. elegans* ACT-2 is most likely present in abundant levels in all cell types. Fluorescence corresponding to the expression of a *GFP::act-2* transgene driven by an *act-2* promoter region is observed in all cells in the early embryo (20 cell stage), in the epidermis as well as in muscle cells^29^. Therefore, we decided to insert a series of genetic modifications identified in the human *ACTB* and *ACTG1* genes in *C. elegans act-2*. In order to span the range of severities observed in human, we chose to study an initial set of eight mutations which included mutations associated with BWCFF (*ACTB* p.L65V, *ACTB* p.G74S, *ACTB* p.G74V, *ACTB* or *ACTG1* p.T120I, *ACTB* p.R196C, *ACTB* p.R196H and *ACTG1* p.T203M)^11,37,38^; as well as non-BWCFF NMA diseases (*ACTB* p.R147S and *ACTB* or *ACTG1* p.R183W) (Figure 1). Among them, *ACTB* p.R196 is a hot-spot identified in over 19 patients, presenting clinical variability with a high degree of brain malformation (83%, Figure 1A)^11,39^. *ACTB* p.R147S represents a hotspot postzygotic disruption causing the mosaic skin disorders known as Becker nevus syndrome^40^. This variant is likely lethal in a homozygous context^41^. *ACTB* p.R183W is associated with dystonia deafness^42–48^.

**Figure 1.**
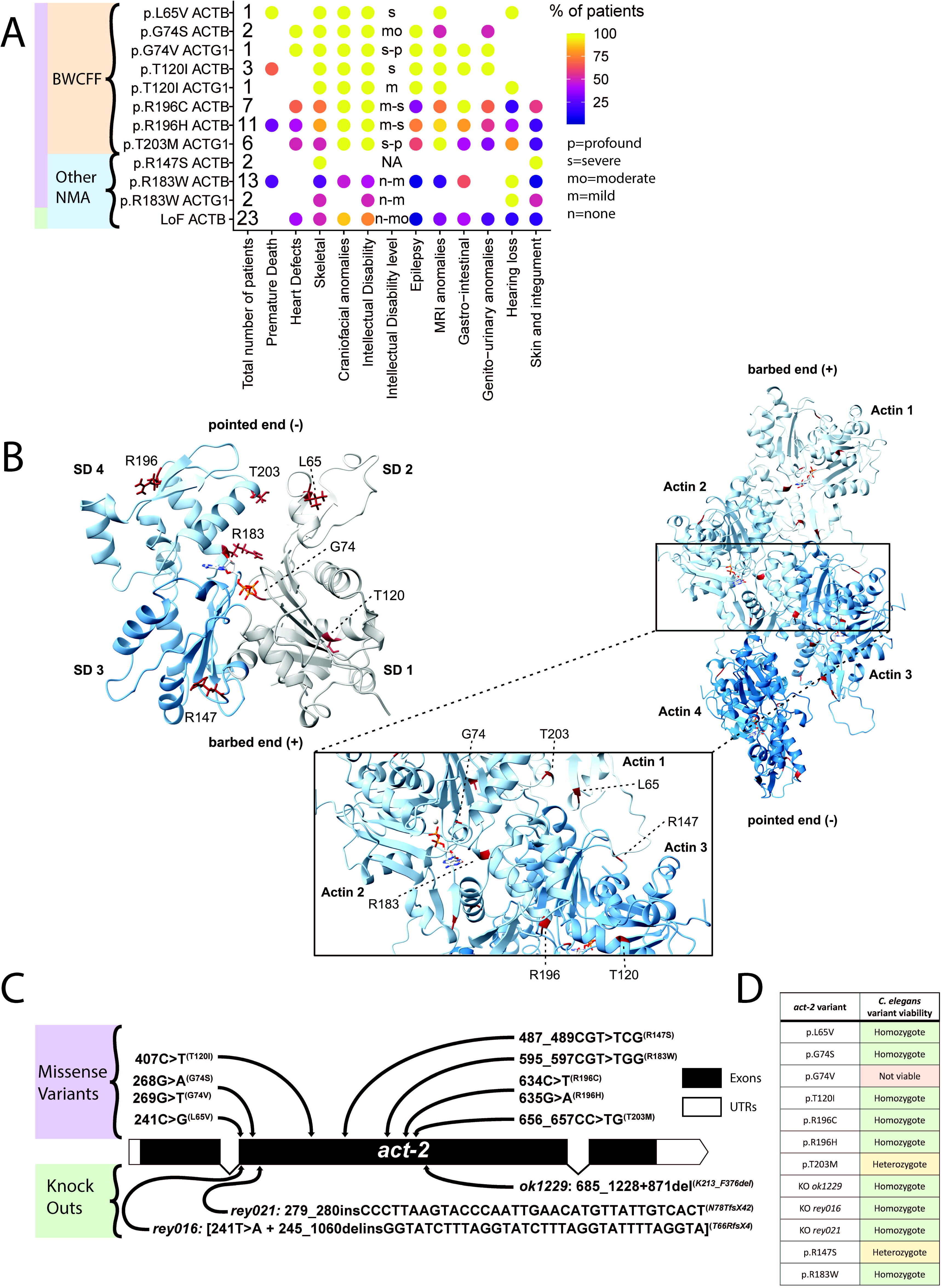
Insertion of Non-Muscle Actinopathies variants in the *C. elegans act-2*. (A) Dot heatmap of patients’ symptoms. Intellectual disability (ID) level letter code : n=none, m=mild, mo=moderate, s=severe, p=profound. MRI = Magnetic Resonance Imaging. BWCFF = Baraitser-Winter Cerebro Fronto Facial syndrome. (B) Structural model of ACT-2 G-actin generated by Alphafold2. Subdomains of ACT-2 are coloured in grey (SD1), light grey (SD2), blue (SD3) and light blue (SD4). (Right) Homology model of the ACT-2 filament (based on PDB-ID: 8DNH). ACT-2 protomers are coloured in different shades of blue. The mutated residues investigated in this study are shown in red. The ATP bound to the individual protomers is shown as a stick-model. The zoom-in shows the location and structural environment of the individual mutated residues in more detail. (C) Schematic *act-2* representation highlighting the location of single point variants obtained via CRISPR edition. Three *act-2* gene knockout alleles are also depicted: *rey016*, *rey021* and *ok1229*^29^. Note that the amino acid numbering is here in frame with human ACTB/ACTG1, consider -1 AA in the frame of *C. elegans* due to an additional cysteine in second N-ter position, the base pair are however kept in the *C. elegans act-2* frame. (D) Table summarizing the overall viability of *C. elegans act-2* variants at 20°C.

We mapped these mutated residues on a structural model of the *C. elegans* ACT-2 monomer (G-actin) and filament to assess the localisation of the variants in different regions of the molecule that correlate with specific functions (Figure 1B). The variants are distributed across all four subdomains (SD) of the actin monomer. Two examined residues (G74, R183) are located within the nucleotide-binding cleft and, although not directly involved in nucleotide coordination or hydrolysis, contribute significantly to the release of inorganic phosphate following ATP hydrolysis^49,50^. The R147 residue is part of the target-binding cleft of the actin monomer, located between SD1 and SD3, which is lined with predominantly hydrophobic residues and represents a main interaction interface for G-actin binding proteins^24,51^. Residue T120 is part of the so-called pathogenic helix of actin (K113-T126), which constitutes a hotspot for mutations in several human actin isoforms leading to BWCFF (cytoskeletal β- and γ-actin), non-syndromic hearing loss (cytoskeletal γ-actin), thoracic aortic aneurysm (smooth muscle α-actin) and nemaline myopathy (skeletal muscle α-actin) (as reviewed in Rubenstein and Wen^52^). Residues R196 and T203 are both located in SD4 and are part of regions of the actin monomer that are heavily involved in the formation of monomer-monomer contacts within the actin filament. The neighbouring residue of R196, E195, forms a salt-bridge with K113 of the laterally opposing monomer. T203 is part of a threonine-rich loop that contributes to lateral and longitudinal contacts in the filament^51,53^. Polymerization deficiencies of the p.R196H and p.T203M variant were verified in *in vitro* experiments. These three variants are predicted to impair actin capacity to polymerise into filaments *in vivo* as confirmed by *in vitro* experiments for the ACTB p.R196H T120I and T203M variants^13,22^. The L65 residue is located in SD2 in close proximity to the DNase-loop that contributes to lateral and longitudinal interfaces and undergoes structural rearrangement upon nucleotide hydrolysis, directly affecting filament stability^49,53^.

Using these selected variants associated with various human symptoms, we generated strains harbouring each NMA mutation in the *act-2* gene, using CRISPR/Cas9 gene editing (Figure 1C, S1A). We verified that no additional mutations were present in the other four actin genes and outcrossed the strains at least four times before analysis. Additionally, we generated two new prospective null alleles for *act-2* (*rey016* and *rey021*) (Figure 1C), each resulting in a premature STOP codon within exon 2 (see methods section for more details). For comparison, we assayed a previously characterized *act-2* knock-out (KO) strain (allele *ok1229*) obtained via an EMS screen, which harbours a larger deletion extending beyond the 3’ UTR^29^.

### ACT-2 missense variants affect the general viability of C. elegans to different degrees

We first compared the general viability of the different mutant *act-2* strains obtained at 20°C. We found that some mutant animals for p.R147S or p.T203M developed into homozygous adults but these few escapers produced very few embryos that were 100% lethal during embryogenesis, suggesting a fully penetrant maternal-effect lethal phenotype. We thus maintained these variants as heterozygous animals using the *nT1* balancer^54^. Interestingly, one mutation (p.G74V) caused lethality in heterozygous state, suggesting a dominant effect for this allele. Indeed, for this variant we obtained very few edited surviving F1 animals after CRISPR injections. Sequencing of several of these F1 edited animals confirmed the presence of a heterozygous G74V variant in the *act-2* gene (Figure S1B); but if some produce very few eggs, none developed into living embryos. Because we were unable to maintain a stable line expressing this variant, we did not pursue its characterization. Surprisingly, another variant affecting the same residue (p.G74S) could be maintained as homozygous. All other variants (p.L65V, p.T120I, p.R183W, p.R196C, p.R196H) as well as the KO *rey016, rey021* and *ok1229* were viable at a homozygous state and were thus maintained as such, although some of them had very few progenies. In conclusion, out of the nine single-point missense variants generated, six could be maintained as homozygous, two as heterozygous and one was inadequately viable (Figure 1D). This initial finding emphasizes how important these variants are in development and health and highlights the remarkable conservation of actin across species.

### act-2 gene knock-out affects reproduction and embryo viability

To assess how individual point variants compare with gene KO, we performed a detailed analysis of phenotypes in the absence of ACT-2 protein expression. We assessed both the newly CRISPR generated *act-2* KO strains (*rey016 and rey021* allele) as well as the EMS generated strain, *ok1229*^29^. We first verified in homozygous L4 animals if transcriptional adaptation of another actin coding gene could compensate for the loss of *act-2*, as proposed by Willis and colleagues^29^. Such transcriptional adaptation has been reported in an *act-5* variant^18,19^. Therefore, we measured the expression of mRNA of all actin coding genes using single animal RT-qPCR in L4 animals devoid of embryos, to avoid developmental differences in the experimental read-out^55^. We found that both *ok1229* and *rey021* strains show a clear absence of *act-2* mRNA and that for *rey021 act-1* expression was increased compared to wild-type (N2) animals (*rey021 act-1* fold change = 2.6; log2fold change = 1.4), unlike other actins which only had milder fluctuations of expression (Figure 2A). We additionally tested whether compensation could be detected in several *act-2* single-point missense variants. We observed some variable mRNA levels of other actin genes than *act-2* only in presence of the variants p.R147S and p.T203M (homozygous L4 assessed), but not for the other variants tested (p.L65V, p.R196C, p.R196H) (Figure 2A). We conclude that there is possibly transcriptional adaptation for actin in *C. elegans*; *act-1* notably compensating for *act-2* loss or for missense variants associated with strong phenotypes (homozygous lethal), as shown in other studies^18,19^.

**Figure 2.**
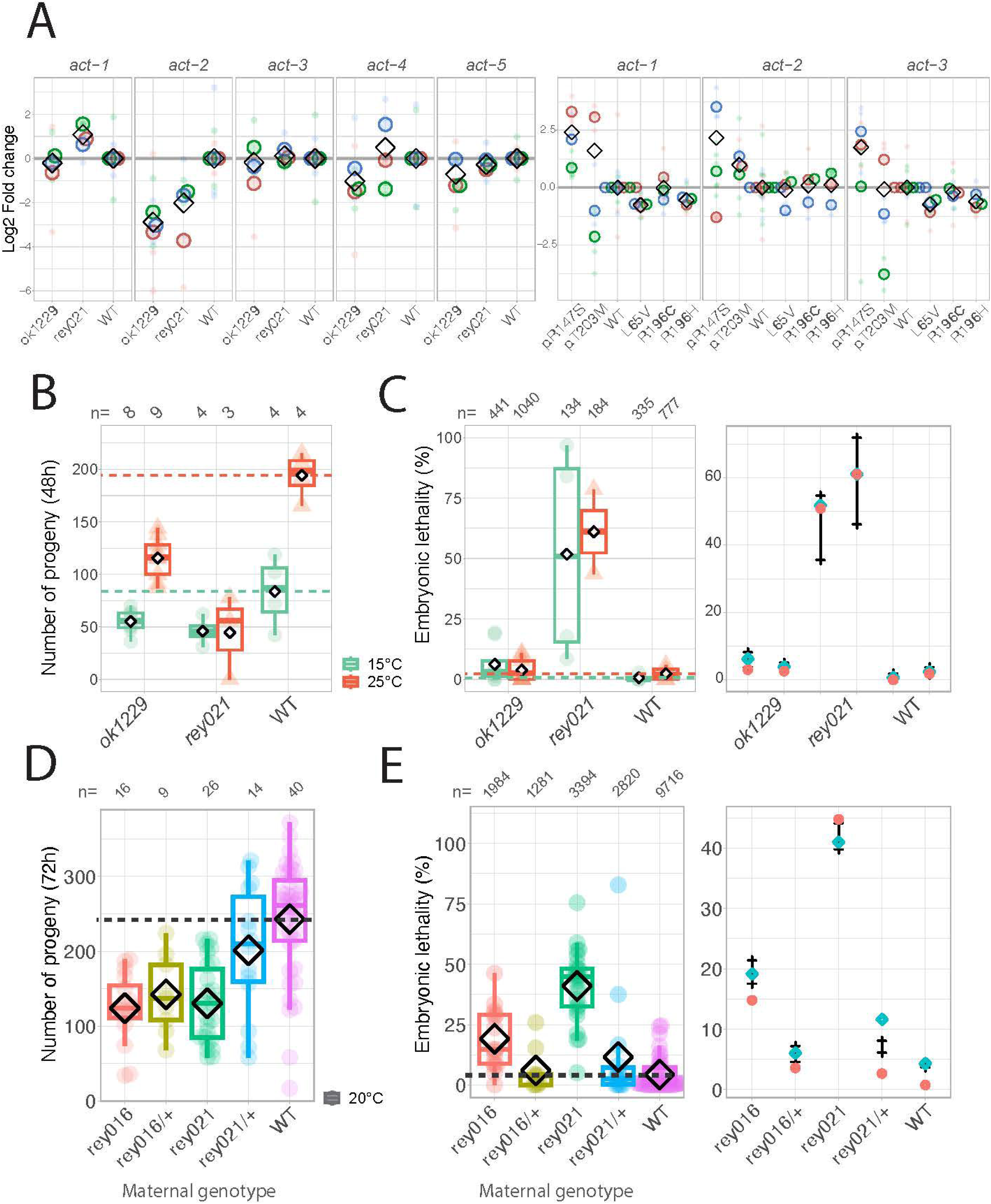
*act-2* gene knock-outs affects fertility despite compensation mechanisms. (A) Single animal RT-qPCR showing the normalized amount of mRNA for each *C. elegans* actin coding genes. Normalization was done using a housekeeping gene (*iscu-1)* and WT values. Different colours show different replicates. Small dot is the value for one animal while bigger dots correspond to the experiment’s mean. Diamonds show the mean of all replicates. (B) Progeny number per homozygous *act-2* knockout (*ok1229* and *rey021*) animal, at the mentioned temperatures, counted during 48h. Standardised boxplots and mean values (diamond) are shown (C) Percentage of eggs that failed to hatch. The mean number of eggs that did not hatch for one animal is shown as a dot or a triangle, overlaid by standardised boxplots and mean values represented by a diamond. The rightmost plot presents the 95% credibility intervals of the posterior distributions of the embryonic lethality rates, considering low informative priors described in the methods section and our observed data set. These are calculated with a Bayesian model based on Poisson laws with an offset to account for the progeny numbers. Experimental values are represented by red diamonds (mean) and cyan rhombus (median). (D) Progeny number at 20°C counted for 72h per homozygous or heterozygous *act-2* KO animal. Standardised boxplots and mean values (diamond) are shown. (E) Percentage of eggs that failed to hatch. The mean number of eggs that did not hatch for one animal is shown as a dot, overlaid by standardised boxplots and mean values represented by a diamond. The rightmost plot presents the 95% credibility intervals of the posterior distributions of the embryonic lethality rates considering low informative priors described in the methods section and our observed data set. These are calculated with a Bayesian model based on Poisson laws with an offset to account for the progeny numbers. Experimental values are represented by red diamonds (mean) and cyan rhombus (median). (B, C, D, E) Boxplots display the maximum and minimum values, the median as well as first and third quartiles.

With respect to fertility and embryonic lethality, the *act-2*(*ok1229)* allele was described as thermosensitive, showing some limited embryonic lethality at 26°C (12/515 embryos, *ie* 2.3 % compared to WT 3/478, *ie* 0.8%) but none observed at 15°C^29^. Therefore, we monitored the brood sizes and percentage of unhatched eggs of *act-2(rey021)* at different temperatures, as well as for WT and *act-2(ok1229)* animals. We observed a reduction in the number of eggs produced by the *act-2*(*ok1229)* and *act-2*(*rey021)* strains compared to the WT reference both at low and high temperatures (15°C and 25°C, Figure 2B). The *rey021* KO allele exhibited a high percentage of unhatched eggs - above 50% - for both temperatures (Figure 2C). However, we did not detect a significant decrease in embryonic development success, in contrast with the previous description of the *act-2(ok1229)* (Figure 2C)^29^. For all tested conditions, we made use of Bayesian methods to infer the 95% credibility intervals of the posterior distributions of the embryonic lethality rates using our data set and both non-informative or informative priors. As shown (Figure 2C), *act-2(rey021)* credibility intervals are clearly separated from that of WT, while this is not the case for *act-2(ok1229)*. Thus, these results established a difference in the developmental success of the different *act-2* KO alleles, the *act-2(ok1229)* allele showing a milder phenotype. As brood sizes and embryonic lethality were similar for the *act-2(rey021)* at both temperatures, we did not see evidence of thermosensitivity for this allele. Thus, all further experiments were performed at the standard 20°C temperature.

To determine if the KO allele effect was dominant or recessive, we quantified brood size and embryonic lethality from heterozygotes hermaphrodites for two CRISPR-generated strains. Overall, the progeny of *act-2* KO animals were reduced compared to WT but *rey021*/+ animals showed a higher progeny number compared to *rey021*/*rey021* (Figure 2D). This effect was not observed for *rey016/+* hermaphrodites, brood size being similar to the homozygous *rey016* condition. Both progenies of heterozygous *rey021/+* and *rey016*/+ hermaphrodites showed an embryonic viability more similar to that of WT (Figure 2E, for example the median embryonic lethality is 3% for the mixed progeny of *rey021*/+ hermaphrodites compared to respectively 45% for *rey021*/*rey021* hermaphrodites and 0.8% for WT hermaphrodites). Taken together, both *rey016* and *rey021 act-2* KO are severely impaired both in terms of fertility and embryonic lethality, this phenotype is only partially suppressed in the heterozygous condition and could therefore be considered as semi-dominant.

### Reproductive capacity is a quantitative readout to classify actin mutants

Assessing general animal fecundity and embryonic fitness in *C. elegans* allows for the quantitative detection of potential defects in oocyte production, embryogenesis or vulva function^56^, three actin-dependent processes. Thus, we measured the number of eggs laid and hatched by single homozygous animals expressing an *act-2* variant at 20°C. A reduction in egg laying was observed for all *act-2* variants, except for the p.G74S and p.R183W variant strains, which appeared unaffected (Figure 3A). The most severe reductions in fecundity were observed for the p.R147S and p.T203M missense variants (mean number of eggs laid: p.R147S = 2, p.T203M = 31, WT = 270), surpassing the reduction seen in the *act-2* KO condition (*rey021* = 78). All other missense variants showed a fecundity reduction less pronounced than the KO animals. Interestingly, heterozygous animals for p.R147S and p.T203M recovered a fecundity closer to WT animals, suggesting an almost complete maternal rescue (Figure S3C).

**Figure 3.**
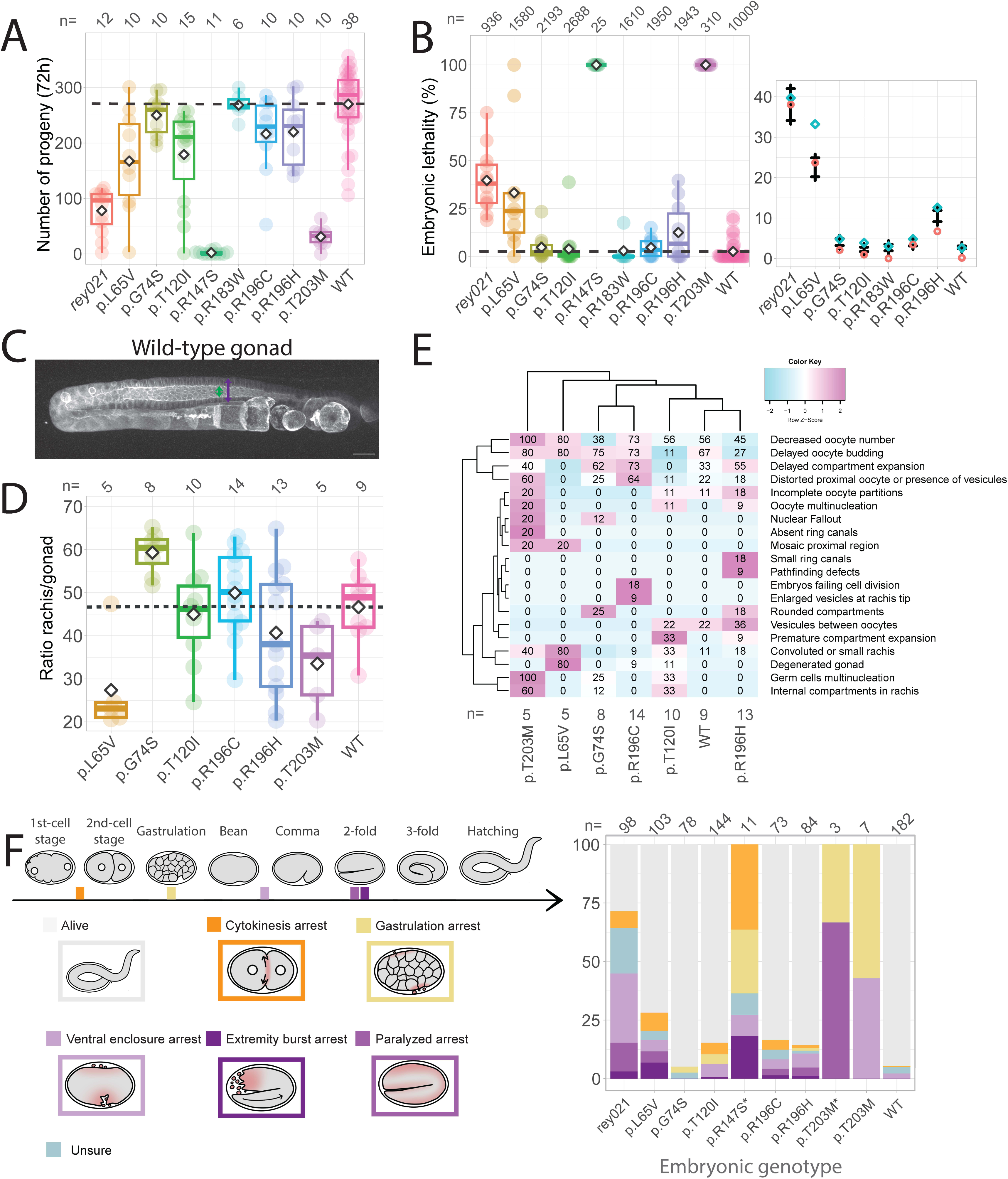
*act-2* variants affect egg-laying capacities and embryogenesis success. (A) Total progeny number at 20°C per homozygous *act-2* variant. The sum of eggs laid for one animal is shown as a dot, overlaid by standardised boxplots and mean values represented by a diamond. (B) Percentage of eggs that failed to hatch. The percent of dead eggs for one animal is shown as a dot, overlaid by standardised boxplots and mean values represented by a diamond. Boxplots display the maximum and minimum values, the median as well as first and third quartiles. The rightmost plot presents the 95% credibility intervals of the posterior distributions of the embryonic lethality rates considering the priors described in the methods section and our observed data set. These are calculated with a Bayesian model based on Poisson laws with an offset to account for the progeny numbers. Experimental values are represented by red diamonds (mean) and blue circles (median). (C) Actin organization in a WT gonad observed using an actin probe (Lifeact::mKate2), max projection of a Z-series of images. Green and purple arrows illustrate the space measured to calculate the rachis/gonad ratio measured in front of the -1 oocyte. Scale bar is 20µm. (D) Boxplot presenting the ratio between the rachis and the gonad width of homozygote animals. Mean is represented as a diamond. (E) Heatmap of observed phenotypes, in percent. (F) Stacked histogram of the end status of embryogenesis for different *act-2* variants in percent. Schemes on the left represent the different defects observed. Also see Videos S1, S2, S3, S4 and S5.

The number of unhatched eggs also varied significantly among actin variants with a trend consistent with that of fecundity (Figure 3B). The p.R147S and p.T203M variants exhibited 100% embryonic lethality when homozygous (Figure 3B), thus displaying a much stronger maternal-effect lethal phenotype than the homozygous *rey021* KO allele (∼41-45% embryonic lethality). Mating with N2 males increased the brood sizes of homozygotes p.R147S and p.T203M without rescuing embryonic lethality (Figure S3A, B). In contrast, p.R147S/+ hermaphrodites showed an embryonic lethality of 17% while that of p.T203M/+ hermaphrodites fell to 3% and for *rey021*/+ to 4% (median values, Figure S3D). This clearly indicates that the p.R147S variant has a more deleterious effect than the null allele, consistent with what has been reported in patients^13^. Note that all the progeny of these heterozygote animals are pooled, thus we here assessed the lethality within a mix of homozygote *act-2* variants, heterozygote and WT embryos.

The p.L65V missense variant also showed significant embryonic lethality (33.2%) although with an increased variability. The p.R196H variant was affected to a significant level unlike the p.R196C variant, as showed by none overlapping 95% credibility intervals of the posterior distributions of the embryonic lethality rates calculated with Bayesian methods (Figure 3B). No other missense variants displayed a significant impairment of embryo survival. Taken together, our data suggest that p.G74S and p.R183W are not impaired in their reproductive capacity; p.R147S and p.T203M display a fully penetrant maternal-effect lethal phenotype; while the other studied variants have much lower penetrance: a mildly decrease in fertility, associated in most cases with embryo viability comparable to WT.

### Gonad internal architecture is impaired in selected mutants

As we observed an impairment in the reproductive capacity of the actin mutant strains, we investigated whether these defects were associated with abnormal organization in the *C. elegans* reproductive system. We performed a detailed analysis of gonad morphology by spinning-disk confocal microscopy using strains expressing both an actin filament probe (mex5p::Lifeact::mKate2), and endogenously labelled non-muscle myosin molecular motors (NMY-2::GFP). *C. elegans* adult hermaphrodites possess two mirror-image symmetric syncytial gonads shaped by highly organized actin structures. These structures include a connected cytoplasm-filled core, called the rachis, surrounded by germ cells with openings facing the rachis and shaped by stable actomyosin rings (Figure 3C). These rings originate from incomplete cytokinesis of primordial germ cells forming the typical specific honeycomb architecture in the functional adult gonad^57^. We first characterized the gonad morphology by measuring gonad and rachis sizes (Figure 3D). In this analysis, the p.L65V variant presented strikingly lower values, whereas the p.T203M showed a more moderate reduction. Interestingly, the variant p.G74S presented higher rachis/gonad ratio than the control even though this is not correlated with a perturbation of fertility. Other variants had values more similar to the WT strain but exhibited greater variances. Second, we quantified defects using established criteria^58^ (Figure 3E). Correlated with the strong decrease of rachis size, the most pronounced phenotypes are found in p.L65V animals, notably illustrated by the presence of 80% of degenerated gonads devoid of oocytes or early embryos. This suggests severe impairment in gonad morphogenesis and/or maintenance in these animals. Still, the penetrance was not 100% as 1 out of 5 p.L65V animals harboured a normal looking gonad, which explains the viability of the strain. Other noticeable phenotypes present in animals expressing different missense variants were: the presence of multinucleated oocytes in 20% of p.T203M and in 11% p.T120I animals; as well as multinucleated germ cells in p.T203M (100%), p.T120I (33%) and p.G74V (25%). Altogether, *act-2* variants are thus associated with abnormal gonad internal organization and multinucleated germ cells, thereby causing a reduction in the number of viable oocytes thus reducing progeny numbers. Lastly, *act-2* could be required for sperm generation or function, as mating WT males to homozygous animals for p.R147S and p.T203M partially rescued the drop of fecundity without improving embryonic viability (Fig S3C, D). To conclude, impaired gametogenesis might explain a reduction in brood size in actin variant genetic backgrounds.

### ACT-2 variants fail at different embryogenesis stages

We next sought to determine causes of the observed embryonic lethality. To this end, we quantified the precise stage at which embryonic lethality occurs in the actin-mutant strains^59,60^ as well as eggshell shape by measuring their aspect ratios and 2D areas (Figure S3C, D). To do so, we monitored the entire duration of embryogenesis using time-lapse transmitted light video-microscopy (Figure 3F). Given the high variability in defects penetrance, high throughput was favoured at the expense of image quality (see Methods). In line with the previous embryonic lethality counts, we detected a variable proportion of *act-2* mutant embryos unable to hatch (Figure 3F). Embryonic defects were categorized into various classes: (1) in the one-cell stage, due to a failure to complete cell division (orange); (2) developmental arrest during gastrulation (yellow), a key event involving cell shape changes and migration, highly dependent on Arp2/3 actin networks^61,62^; (3) developmental arrest at later steps of development (three shades of purple). Class (3) was including most cases of defects and can be further subdivided into: (3i) arrest during ventral enclosure (light purple), a process also known to rely on both protrusive and contractile actin-related events^63^; (3ii) arrest during elongation, also referred to as Pat embryos (for paralyzed at two-fold, purple), suggesting muscle-defective embryos and/or defective hemidesmosomes; and (3iii) rupturing embryos during elongation, an obvious sign of impairment of tissue mechanics (dark purple)^64^. Interestingly, the total percentage of defects and their frequencies varied among the different genotypes but the KO condition showed a high proportion of lethality in each. In contrast, homozygous p.R147S embryos suffered severely from early embryonic death (cytokinesis and gastrulation defects) while for heterozygous p.R147S embryos, early death was reduced and they tended to die more frequently during ventral enclosure or in later stages of elongation. As elongation requires efficient muscle contractions, this result corroborated the p.R147S impaired muscular capacity measured in the motility assay (Figure 5A, see below). The few embryos expressing the p.T203M allele, arrested at different stages of morphogenesis between gastrulation and elongation with different ratios depending on the genetic condition. These findings suggest that the p.R147S and p.T203M, both in homozygote and heterozygote embryos, are more detrimental or toxic for early embryogenesis than the complete or partial absence of ACT-2 protein (*rey021* condition), supporting a presumptive semi-dominant effect. To conclude, this analysis revealed variant-specific differences in the mechanisms leading to actin-related lethality, likely associated with fine-tuning of specific actin organization, dynamics or mechanoproperties in key stages of embryogenesis.

**Figure 5.**
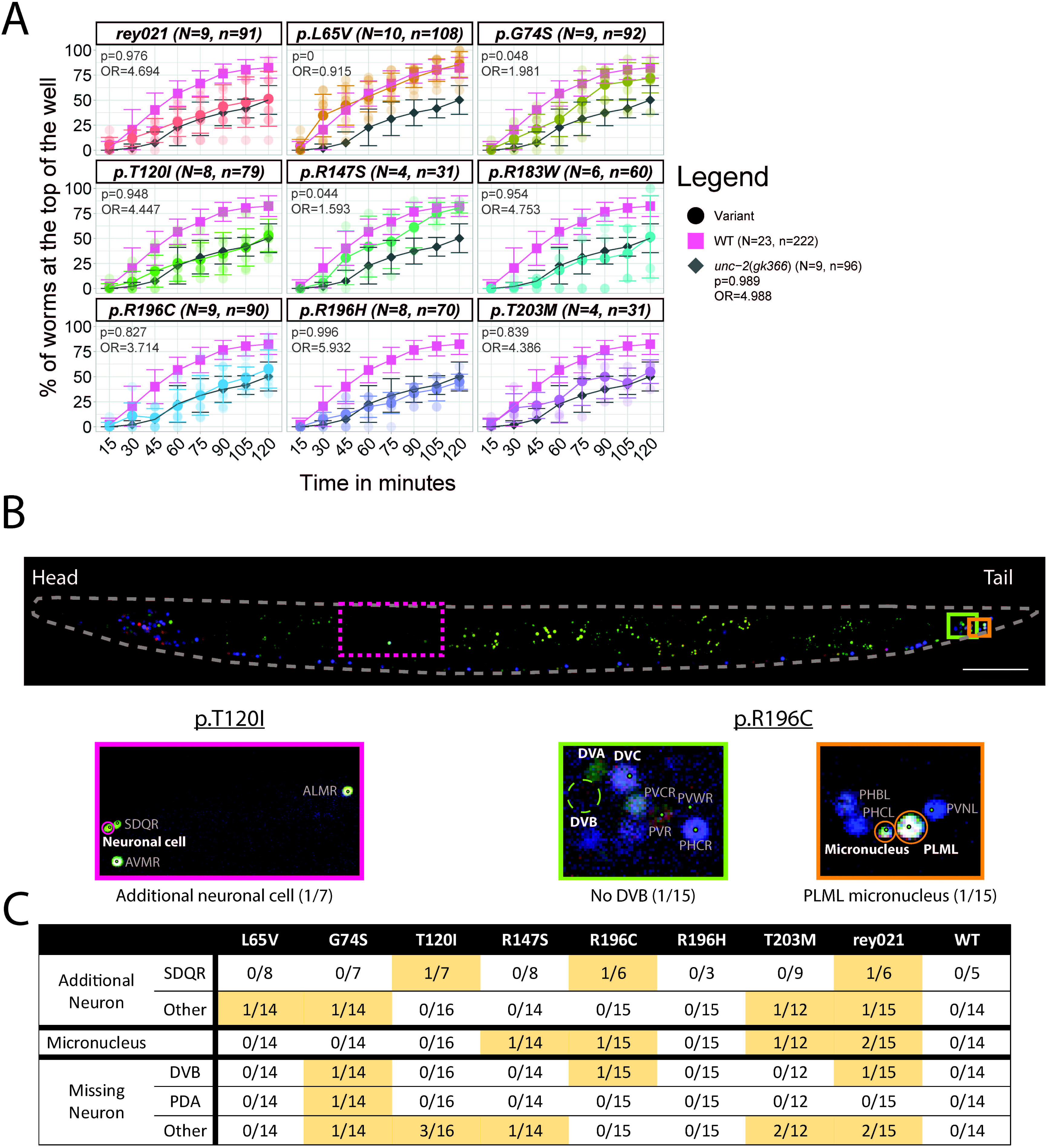
Actin mutations can affect motility and/or neuronal cell number. (A) 3D animal migration assay. Percentage of animals reaching a chemoattractant after migration inside a pluronic gel, monitored every 15 minutes for 2 hours. Results for individual wells (N) for each variant are visible as transparent dots, n is the total number of animals assessed and bars show the standard deviation. The final proportion for each variant was compared to that of WT in the frame of Bayesian statistics, using two binomial distributions. The odds ratio (OR) as well as the probability (p) that the difference is superior to 0.2 are indicated. (B) Variation of the neuronal cell number are observed in *act-2* variant conditions. Reconstructed image of an animal expressing the NeuroPal transgene. Insets show representative images of identified defects compared to the WT *act-2* reference: presence of a micronucleus (Orange), an extra (Magenta) or a missing cell (Green). Scale bar is 50 µm. (C) Summary of neuronal phenotypes observed in the actin variants.

### Altered cell mechanics but surprisingly mild cortical actin disorganization in early embryos of several act-2 variants

We then asked whether defects emanate from abnormal cytoskeletal organization or impaired force production, such as abnormal cytoskeletal contractility linked to actin-myosin interaction. These possibilities are not mutually exclusive and could be linked, as architectural defects can impair contractility^65^. We thus increased the time and scale resolution by performing spinning-disk confocal microscopy on strains carrying the Lifeact probe to follow actin filament organization as well as an endogenously labelled non-muscle myosin (NMY-2::GFP) to assess contractility during the first cell divisions of *C. elegans*^66^.

With respect to cortical architecture and dynamics, we were surprised by the normal general organization of the actin networks observed in some *act-2* variants (Figure 4A), even though reduced polymerization capacities have been quantified biochemically using purified proteins as for p.R196H^22^ or p.T120I and p.T203M^13^. Visual inspection did not reveal major signs of impaired actin density or general organization in most embryos. For a few embryos, we did observe lower density regions or holes in the cortex as for some p.T203M and p.L65V embryos (Figure 4A). Similarly, we did not detect signs of impairment in the reproducible sequence of events taking place in zygotes for most embryos. These events, which highly depend on proper cortical organization and contractility, include polarity establishment associated with cortical flows, pseudo-cleavage formation, and the maintenance phase where polarity is stably maintained^67,68^. This was evidenced by the unchanged size of the AB daughter cell in *act-2* variant conditions (Figure S4). It is possible that many NMA actin variants can copolymerize with WT actin, resulting in mild phenotypes of disorganization that are still compatible with life in both humans and *C. elegans*.

**Figure 4.**
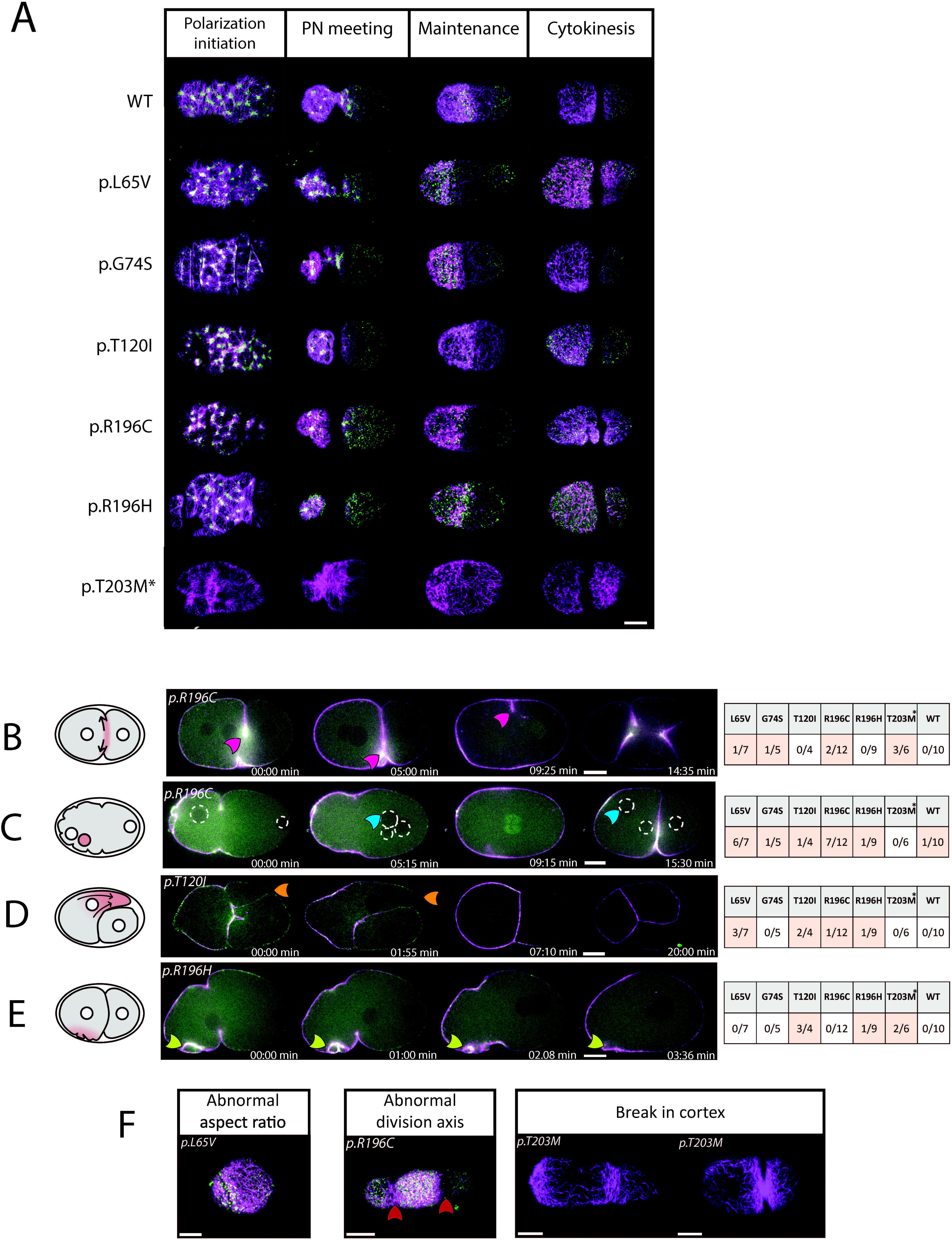
Actin mutation effects on cortical organization in early embryos. (A) Cortical plane of embryos at different stages, Lifeact::mKate2 shown in magenta and NMY-2::GFP shown in green, to reveal actin filaments and myosin distribution. p.T203M animals only have Lifeact::mKate2. (PN = Pronuclei) Anterior is on the left. (B, C, D) Equatorial plane image of embryos expressing Lifeact::mKate2 (magenta) and NMY-2::GFP (green). (B) Cytokinesis defect observed. The cytokinesis ring relaxes after completion of a first ingression (pink arrow), this is followed by multiple simultaneous ingressions as the cell proceeds to divide directly into four daughter cells. See Video S6. (C) An additional nucleus is visible (blue arrow). Nuclei are circled. See Video S7. (D) Blebbing. Large protrusion called bleb extrude from the anterior cell (orange arrow). After actin and myosin reassembly, the bleb retracted and division seemed to occur normally. See Video S8. (E) Abnormal membrane ruffling is observed in some embryos (white arrow). (F) Cortical plane of embryos presenting other phenotypes. Abnormal aspect ratio of the cortex implies a rounder embryo (see also Figure S4). Abnormal division axis (red arrows) leads to abnormal cell-cell contacts. Breaks in the cortex are highlighted by green arrows. (*) Embryos from homozygote actin variants expressing Lifeact::mKate2 only. Scale bar is 10µm.

Although most cells maintain seemingly normal actin cortical networks, we observed some cell-scale defects, revealing impaired cytoskeletal properties in some embryos (Figure 4B, C, D, E, F). As expected from previous analyses, we identified cell-division defects where ingression progresses smoothly but subsequently retracts (Figure 4B). This suggests that cytokinesis-ring assembly and contractility might not be impaired, but rather that failure happens at the last steps of cytokinesis completion during abscission. We also observed a supernumerary nucleus present either in the AB cell or in the P0 zygote, possibly caused by cytokinesis failure in germ cells or meiosis defects due to incomplete polar body extrusion in the oocyte (Figure 4C). As meiosis and cytokinesis completion share common mechanical principles linked to actin, such as the formation of a contractile actin ring required to bring together closing membrane domains^69,70^, these failures may share a similar origin. In addition, we noticed the presence of blebs in some embryos (Figure 4D, E), note that blebs were also reported for two previously described *act-2* missense variants the *or621* and *or295* alleles^29^. Blebs are transient cell protrusions that can be triggered by two main events: the detachment of the cortex from the plasma membrane or the formation of a crack in the cortex. Blebs then grow due to the internal hydrostatic pressure imposed by cortical contractility^71^. Blebs were mostly observed at the 2-cell stage, at the interface between the AB and P1 cells before cell rounding prior to division. We observed larger blebs, mostly in p.L65V, p.T120I and p.T203M variants, emanating from AB, sometimes dragging the nucleus into the bleb, before a fast retraction period. Retraction coincided with the increase of myosin at the cortex at cytokinesis onset. Embryos presenting the blebbing phenotype also had transiently abnormal cell shapes, an overall less round, wavier cortical surface (Figure 4D) and seemingly diminished cell-cell contacts, particularly between P1 and ABp cells at the 4-cell stage. This phenotype resembles cortical blebs that were proposed to reposition the cytokinesis ring of the zygote in *ani-1(RNAi);pig-1(gm344)* embryos in the presence of excessive myosin accumulation at the anterior cortex^72^. While we did not observe mispositioning of the furrow or abnormal nuclei migration, the similar phenotype suggests a shared site of cortical instability.

### Motility is mildly impaired for a few act-2 variants

Like its actin cytoplasmic paralogues, ACT-2 protein is also required in muscle cells^29^, evidenced by dominant mutations in *act-2* impacting *C. elegans* motility, with uncoordinated (Unc) phenotypes on 2D surfaces^32^. As no such strong Unc phenotype was observed in the studied *act-2* mutants, we quantified their capacity to migrate inside a pluronic gel towards a chemoattractant (food source) in a 3D burrowing assay^73^. We observed that 82.4% of WT animals and 50.2% of mobility-deficient control animals (*unc-2*(*gk366*)) reached the food after 2 hours (Figure 5A). We detected altered 3D migration for the KO *rey021* and for the variants p.T120I, p.R183W, p.R196C, p.R196H and p.T203M, while other variants showed unaffected migration (p.L65V, p.G74S, p.R147S). We verified that these effects were not due to impaired chemosensation (Figure S5). Note that in this assay we observed an increased number of immobile animals for variants *rey021* and p.T203M only. In conclusion, six out of nine assessed *act-2* variants have impaired 3D motility, while two variants display a substantial percentage of animals that are defective both for 2D and 3D movement.

### Low penetrance of neurogenesis defects

In humans, neurodevelopmental defects are the most frequent symptoms across the NMA spectrum. We thus studied neurons in *C. elegans* L4 animals. To this aim, we took advantage of a multicolour neuronal labelling transgene present in the NeuroPAL strain^74^. The NeuroPAL transgene codes for the expression of a combination of 41 neuron specific reporters for gene expression and neuronal activity, associated with four different fluorophores. This labelling allows each neuron in the animal to be identified by a unique set of fluorophores, distinguishing them from neighbouring neurons. After introducing the NeuroPAL transgene in the actin variant strains, we mapped all neurons present in the animal’s midbody and tail, which include every type of neuron. For every observed animal, we did not detect altered neuronal positions. However, we did observe rare events, for each allele except p.R196H, wherein we detected the presence of a supernumerary neuron (adjacent to SDQR or in the lumbar ganglion) or a missing neuron (such as DVB or PDA, Figure 5C). These altered numbers suggest that NMA neurodevelopment defects may be reproduced in our mutant models.

### Data integration to build a comparative tool

Taking advantage of the multiscale characterization performed on each mutant, our final goal was to classify the different missense variants in terms of severity. We used each of the quantitative results as variables to create a Principal Component Analysis (PCA) graphical representation (Figure 6, variables available in Figure S6). This dimension reduction allowed us to group phenotypes into different classes and allowed us to identify the few most promising variables to be used for classification purposes (Figure S6). In the PCA plot, the WT strain was located in the left side close to the mildly affected variants (p.G74S, p.T120I, p.R183W, p.R196C, p.R196H), thus defining class I: characterized by negative values along dimension 1. One can note the singular position of the p.L65V, isolated at the boundary of the class I group, emphasizing some specificity with respects to this variant. This result correlated with human data, where an *ACTB* p.L65V patient presented severe symptoms and early death. The KO *rey021* strain was also isolated in a more central position. Stronger phenotypes such as in p.R147S or p.T203M variants were positioned in the right half, defining class II (positive values along dimension 1). Euclidian distances calculated between each variant and the wild type reference confirmed the choice of groups: class I variants were grouped bellow a distance of 3 and class II positioned above this threshold (Figure 6B). Interestingly, heterozygous variants (*rey021*/+, p.R147S/*+* and p.T203M/*+)* were positioned closer to the WT than their homozygous counterparts, thus in accordance with a reduction of the severity of the defects in presence of one wild-type *act-2* allele. Differences in the directionality of these repositioning trajectories probably underly differences in the mechanism at the origin of the defects that may affect differently the scored phenotypes (Figure S6A). To assess the robustness of our analysis we performed additional PCA either removing all the data linked to the insertion of fluorescence in the *act-2* variant genetic background, thus keeping the most easily quantifiable phenotypes (Figure S6C, D) or removing the two most lethal strains p.R147S and p.T203M (Figure S6E, F). The similarity of the PCA 6B and S6C showed that the removal of these specific variables did not modify the overall positioning of strains, whereas the PCA S6E confirms the respective positioning of the homozygous variants for example, p.L65V notably clearly apart from WT and other *act-2* variant studied. In conclusion, using quantitative multiscale analysis of a panel of actin variants, we have provided access to a valuable comparative tool enabling the positioning of newly identified variants on a graded map of survival potential, if such new variants were obtained and analysed using the same parameters reflected in our PCA analysis.

**Figure 6.**
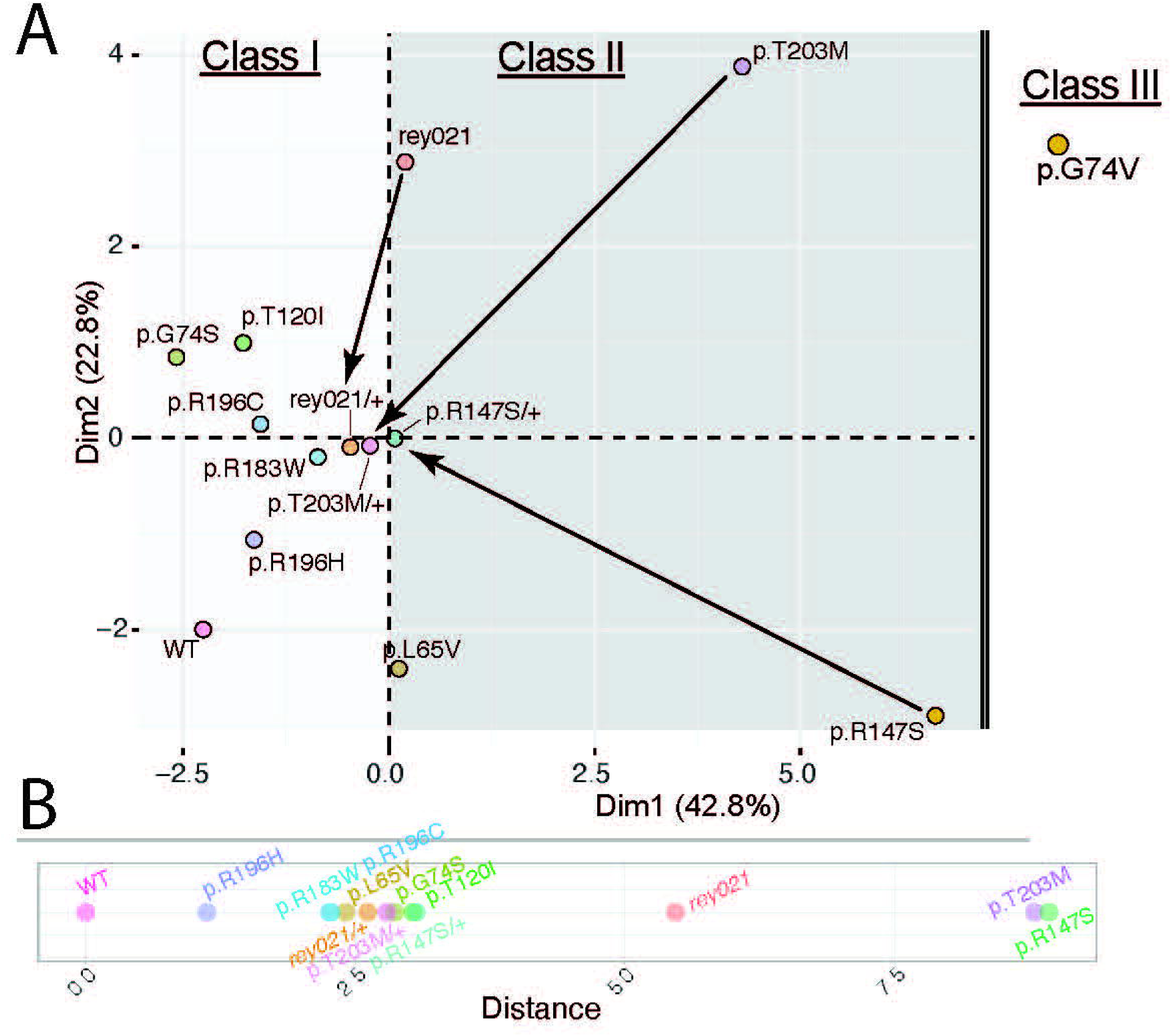
Data integration to build a comparative tool. (A) Principal Component Analysis plot of analysed actin variant animals reveal clusters which can be associated to classes of phenotypes. Corresponding table with data is available in Figure S6A. p.G74V was added manually as no data could be retrieved for this strain. Arrows link homozygotes to their respective heterozygotes. (B) Distances calculated between each actin variant and the WT reference in the two first dimensions.

## DISCUSSION

Patients affected by the rare disorders from the Non-Muscle Actinopathies (NMA) spectrum and its subclass BWCFF, exhibit a large range of symptoms and diverse survival potential, leading to complicated medical prognostics. Given the ubiquitous nature of actin, which is involved in numerous cellular events at all stages of development and throughout lifespan, it is important to analyse multiple phenotypic parameters in animal models. Ideally in order to effectively distinguish the impact of each actin variant, these parameters should span diverse length scales: from macroscopic actin organisation to cell behaviours and phenotypes at the organism scale. In this study, we used the fast-growing and simple model organism *C. elegans* to study *in vivo* the quantitative impact of some of the associated single cytoplasmic actin variants. Using CRISPR/Cas9 genome editing, we successfully generated new *C. elegans* models expressing endogenously either *de novo* actin variants identified in human patients or premature STOP mutations to reproduce loss-of-function conditions. For each variant, we quantified general fitness, reproductive capacities, muscle-based motility, embryonic development, and neuronal differentiation. The final step using an integrated PCA analysis enabled us to summarize the phenotypic classification, providing a valuable tool to position newly identified variants of unknown significance on an existing map. We foresee that these data support the use of *C. elegans act-2* mutations as a pertinent and promising model for the investigation of NMA.

A key finding of the functional characterization of *act-2*, was the significant phenotypic difference between different *act-2* knock-out strains *(rey021, rey016, ok1229).* In summary no *act-2* mRNA was detected for allele *act-2(rey021) and act-2(ok1229)* but an upregulation of *act-1* mRNA levels was clearly observed only for *act-2(rey021)*. Regarding fertility, brood sizes were reduced in all homozygous KO conditions but were variable in the heterozygote *rey021/+* and rey016/+ conditions (partially improved only in the *rey021/+* condition). Regarding embryonic lethality, the phenotypic differences between alleles were even more pronounced: it varied from no embryonic lethality for *act-2(ok1229)* to different penetrance of lethality for *rey016* and *rey021* that were strongly but not fully rescued in heterozygote maternal conditions. Therefore, the *act-2(rey021)* and *act-2(rey016)* alleles showed a semi-dominant effect as they are not fully rescued in the heterozygote condition. To conclude, we suspect that these alleles are haplo-insufficient, with both a maternal and zygotic effect, unlike *act-2(ok1229)* that was associated with reduced brood sizes but presented a comparable embryonic lethality as wild-type. Such a difference among the KO strains was unexpected and we can only hypothesize on its origin: it could probably be linked to the presence of different and intricate mechanisms allowing to sustain actin homeostasis compensating its loss. These mechanisms include genetic compensation also referred to as isoactin switching, dosage compensation and transcriptional adaptation. Additionally, the presence of different feedback mechanisms either relying on the gene locus, mRNA, or the actin protein itself, may be at work differentially in these genetic contexts^26^. The absence of major changes in other actin expression levels for some missense variants (L65V, R196H, R196H) but presence of changes for more deleterious missense variants (R147S, T203M) reinforces the possibility that different levels of regulation may be at play. Note that different compensation efficiencies may vary both between embryo and L4 animals and may have some tissue specificity.

For *ok1229*, it is also possible that a linked non-identified second-site modifier (eg. a linked suppressor mutation) was introduced during non-targeted mutagenesis, resulting in the observed differences with the CRISPR-generated alleles. As in human where loss of a cytoplasmic actin allele induces a compensatory transcriptional adaptation^7,24,25,75^, *C. elegans* showed transcriptional adaptation to compensate for *act-2* loss, as the closely related actin paralogue gene *act-1* was found to be upregulated in the *rey021* condition. One hypothesis for the absence of compensation in other conditions, could be the loss of a 25bp key sequence located in *act-2* second intron that was shown to both regulate actin genes expression and be required for transcriptional adaptation^18,19^, or loss of another regulatory element in the deleted regions of *ok1229* or *rey016*. This sequence is indeed removed both in *rey016* and *ok1229* but not in *rey021* in which it may still act to induce an overexpression of *act-1* via mRNA-linked transcriptional adaptation. Note that no significative change was observed in *act-3* levels in these contexts.

Beyond regulation via paralogue gene expression, actin levels are additionally regulated at posttranscriptional and posttranslational levels and actin concentration was also shown to regulate actin mRNA synthesis, all these effects are possibly differentially affecting cytoplasmic actin concentrations in these genetic contexts^76,77^. Overall, this shows that a fine tuning of the actin abundance is at work and impacting the different function we assessed, and that the presence of ACT-2 is notably important for maintaining a high fertility and ensuring the success of embryonic development.

Another significant finding was that three actin variants exhibited stronger phenotypes than the *rey021 act-2* KO, namely p.G74V, p.T203M and p.R147S. The p.G74V variant led to animals producing 100% lethal embryos even when heterozygote, therefore harbouring a strong dominant maternal-effect lethal phenotype. In contrast, embryos from variants p.T203M and p.R147S were 100% lethal only when the mother was homozygous, showing a fully penetrant maternal-effect lethal phenotype, with a stronger effect than that of null alleles. However, the latter mutations were strongly improved in heterozygotes animals to a comparable extent than the *rey021/+* condition. Still, especially for the p.R147S missense variant, heterozygous animals displayed a low embryonic lethality consistent with a semi-dominant allelic effect. This could indicate that the presence of this mutated actin residue disrupts proper cytoskeletal function, either through essential perturbations of actin filament assembly or by interactions with actin-binding proteins. These two variants p.G74V, p.R147S and possibly p.T203M could act as semi-dominant negative alleles by antagonising normal polymerisation kinetics of the wild-type actin and thus affecting a diversity of cellular function notably required for the success of embryonic development. To distinguish dominant negative effect from other gain-of-function mechanisms, one could test if increasing amount of wild-type protein is enough to rescue the lethality. At the biochemistry level, a simplification considering actin polymerisation kinetics in isolation, such hypothesis could be easily tested using a range of protein concentrations^20,22,50^. However, actin polymerisation is just one of the many parameters of actin *in vivo* role and is probably not sufficient to explain phenotypes of more complex *in vivo* situations. Indeed, if both p.R196H and p.T203M have impaired *in vitro* polymerisation rates, only p.T203M allele is severely impacting *C. elegans* fitness and embryonic success. Thus, the combination of both findings, *in vitro* biochemistry and *in vivo* animal models, provides key information to determine the impact of each actin variant.

Using these models, we observed an unexpected striking difference between two substitutions at the same residue. Animals expressing the p.G74S variant show only mild effects on embryonic viability whereas the p.G74V animals are significantly more affected, as mentioned above. This position is of particular interest because of the neighbouring His73 which is known to undergo irreversible methylation, regulating actin’s interdomain flexibility in multiple species and known to affect smooth muscle contraction in mice for example^50,78^. Additionally, due to its location within the nucleotide-binding cleft, this variant might impact the phosphate release following actin monomer incorporation into the filament. Consequently, the binding of ADF/cofilin would be perturbed, leading to impaired disassembly kinetics. Of note, *C. elegans’* ADF/cofilin homologue UNC-60 induces lethality when knocked down^79^. Biochemical analysis and *in vitro* studies using purified actin variants could examine these hypotheses and compare biochemically how the presence of the hydrophobic Valine is more detrimental than Serine.

Additionally, and from a more fundamental perspective, characterizing actin variants is also powerful to reveal new properties of actin networks. The observed abnormal internal gonad organization is highlighting the importance of ACT-2 for proper cytoskeleton architecture establishment in this tissue. The presence of multinucleated cells revealed the importance of ACT-2 in cytokinesis and possibly meiosis. Blebs, on the other hand, are intrinsically linked to cortical contractility and to cortex-to-membrane attachment, thus reinforcing the possibility that membrane-to-cortex attachment might be impaired, as also suggested by late cytokinesis defects. We use this rationale to propose that variant p.L65V is detrimental to gonad cytoskeleton organisation and functional oocyte production; p.R147S possibly impairs mostly branched actin networks essential at different morphogenesis stages; and p.T203M, p.T120I, and p.R196C/H impact contractility as indicated by blebbing, paralysed arrest and impaired motility. As we have demonstrated that developmental arrest at critical embryonic stages occurs at varying frequencies depending on the nature of the variant or copy number —in the case of p.R147S— we propose that these key developmental stages act as specific “barriers” to developmental success, barriers of different difficulty linked to the precise site of mutation, that must be successively overcome during embryogenesis. As developmental timings^80^ were not drastically affected (Figure S3G), we suspect that these barriers might have a biomechanical origin. Alternatively, it is possible that some stochastic defects in the early embryo such as lineage defects or mispositioning of cells could lead to developmental arrest at later cell stages. A recent study by Xiao and colleagues^81^ sheds light on how cellular plasticity can buffer genetically-induced defects in *C. elegans* embryos. Thus, another hypothesis would be that our variants do not only impair morphogenesis per say - at the observed timing of arrest-but might be resulting from earlier defects which could have been initially compensated. A more detailed analysis of early embryonic development, such as using automated profiling of embryonic development^82,83^ or lineage tracing methods^84^ could enable to verify such hypothesis.

Observed defects commonly display incomplete penetrance and the general organization of actin at a macroscopic scale is often remarkably unaffected. This suggests an adaptation of cytoskeletal actin to perturbations, possibly originating from the cell’s ability to maintain the adequate large-scale physical properties of the cortex, also known as “morphogenetic degeneracies in the actomyosin cortex”^85^. Compensation mechanisms may ensure the robustness of embryonic development by maintaining proper cytoskeleton mechanical properties. Note that a similar robustness to cytoskeletal perturbation, in that case a resistance to microtubule disassembly, was observed in the context of *C. elegans* embryonic elongation^86,87^. In the context of actin, this could also be explained by a functional compensation by other actin isotypes expressed in early embryogenesis, notably ACT-1 or ACT-3, which might compensate for polymerization defects. *In vitro* experiments with purified proteins using mixtures of WT and mutant actin, such as p.R196H, also show a partial rescue of polymerization defects^22^. Thus, biological variations such as gene expression fluctuation, gene compensation, actin and ABP concentrations, mechanochemical feedback or fluctuation of properties impacting cell or tissue mechanics, determine the success of embryogenesis for each individual. Consequently, each embryo may respond differently to these compensation mechanisms and might arrest at a different developmental stage. In order to go a step further, a detailed analysis of actin spatio - temporal organisation and actin molecular dynamics together with the study of some key actin binding proteins such as myosin, formin or capping-protein both in the early embryo as well as during some key stages of development, will enable to reveal additional details on the respective molecular effects of these actin variants in a living animal.

Finally, the initial investigation of the consequences of actin perturbation on neurogenesis revealed a defect in acquiring the correct number of neurons, albeit not penetrant. SDQ neurons are born from an asymmetric division followed by the death of their sister cell while DVB and PDA are born from the trans-differentiation of a rectal epithelial cell^88^. Even at low penetrance, these numbers are notable, due to the 100% invariance of these neuronal lineages in *C. elegans* under wild-type conditions^88,89^. These findings suggest a potential defect in the process of neuronal asymmetric division or fate acquisition, which might be amplified in the context of human cortical development where neuron numbers are far greater and asymmetric divisions have been reported in the striatum. The asymmetric divisions required for neuronal fate acquisition can depend on the asymmetry of Arp2/3 nucleated cortical actin polymerisation^89^. As both actin polymerisation and Arp2/3 nucleation were already described to be impaired for some variants studied *in vitro*^22^, this result is not surprising and will require further investigation in *C. elegans*. Such additional studies may lead to advances in identifying the links between the molecular and functional levels that underlie NMA neuronal pathophysiology. Importantly, the NeuroPAL data using nuclei as proxy of neuronal cell positioning, did not reveal other gross defects in terms of cell migration. This might be due to the redundancy at work during development that ensures the robustness of final neuronal positioning in *C. elegans*. Assessing migration dynamics and axonal networks may uncover additional abnormalities not visible in the current analysis.

As all animal models, the use of *act-2* variants in *C. elegans* has some limitation. First of all, we do not have a confirmed correspondence between the three *C. elegans* cytoplasmic actin paralogues and the human *ACTB* or *ACTG1* genes. Even if ubiquitous, the *act-2* expression may vary in abundance in *C. elegans* cells or tissues compared to the two human cytoplasmic actin, thus introducing possible bias or misinterpretations. For instance, the key role of ACT-2 for proper gonad organization and fertility might be specific to *C. elegans* and might not have a direct equivalent in human. We however think that the highly conserved *in vivo* actin-related biochemistry is ensuring, notably in the case of strong perturbation, that variants identified as inducing strong perturbation in *C. elegans* also have a negative impact in human cells.

Altogether, this work provides a new animal-model library for the study of actin variants associated with NMA. We show that different non-synonymous mutations in nematodes lead to a broad range of phenotypes as in humans, and that the severity of *C. elegans* phenotypes correlates with patients’ survival potential. For example, two severely affected *C. elegans* strains are p.R147S and p.T203M. This correlates with the observation that several *ACTB* p.R147S patients were identified harbouring only post-zygotic mutations, thus expressed as mosaics in a subset of epithelial cells suggesting a lethal issue in human if expressed in all cells^40^. Additionally, several patients with variant p.T203M in *ACTG1* suffer from severe BWCFF symptoms including microcephaly, profound intellectual disability and non-ambulance^13^. This residue is known to be a hot spot for *ACTG1* but no patient with an *ACTB* mutation has been identified so far, raising the question of lethality for *ACTB* in humans. Similarly, the *C. elegans* variant p.G74V showed 100% lethality and patients with p.G74V variant in *ACTG1* are severely affected, while no *ACTB* p.G74V variant has been discovered so far. To conclude, the severity of this initial set of *C. elegans* mutants correlates with that reported in human patients.

With commercial gene editing services in *C. elegans* offering quick delivery times and automated analysis of various criteria, it is plausible that *C. elegans* NMA models may be used in the future as indicators to aid clinical prognosis for humans. Additionally, all the tools generated here could be used to investigate further the mechanisms underlying NMA pathophysiology, including genetic suppressor screens and/or drug screening for therapeutic strategies.

### Limitations of the study

In our opinion, this study has three major limitations. As the correspondence between the *C. elegans* cytoplasmic actin orthologues and the human ACTB or ACTG1 genes is still unknown, we used *act-2* to model variants of both types. All three proteins are highly similar and the most probable cause of the identified functional differences between ACTB and ACTG1 possibly lies in their genetic encoding^15,23^. Consequently, even if ubiquitous, *act-2* expression may vary in abundance in *C. elegans* cells or tissues compared to the two human cytoplasmic actins, thus introducing possible bias or misinterpretations. Second, we report strikingly different phenotypes for different null alleles in the *C. elegans act-2* gene: the *ok1229* allele showed a very moderate phenotype in contrast to the *rey021* or *rey016* alleles. Although null alleles are expected to induce similar phenotypes, several possibilities may explain this discrepancy, notably the presence of variable compensation of actin proteins (also see the discussion). In addition to the observed rescue of embryonic lethality in heterozygotes (*rey021/+* or *rey016/+),* to further address this issue, a rescue experiment with a wild-type version of an *act-2* transgene could have been performed to rule out the existence of potential linked mutation(s) causing the phenotype. Third, concerning gene compensation between actin genes, we must admit that our RT-qPCR experiments showed a high variability, likely arising from a biological source as technical replicates were more alike. Given that these experiments were performed on individual animals, it is possible that this variability may come from differences in the level of genetic compensation at work in the selected animals.

## RESOURCE AVAILABILITY

### Lead contact

Requests for further information and resources should be directed to and will be fulfilled by the lead contact, Anne-Cécile Reymann.

### Materials availability

All generated *C. elegans* strains generated in this study are available from the lead contact -if not already available at the *Caenorhabditis* Genetics Center (CGC).

### Data and code availability

Data: All data reported in this paper will be shared by the lead contact upon request.

Code: All original code has been deposited and is publicly available via our GitLab (https://gitlab.com/igbmc/reymannlab/2025-hecquet-iscience).

## Additional information

Any additional information required to reanalyze the data reported in this paper is available from the lead contact upon request.

## ACKNOWLEDGMENTS

This work was funded by European Joint Program for Rare Diseases through the European Union’s Horizon 2020 research and innovation program under the EJP RD COFUND–EJP N° 825575. N.D.D. and D.J.M./J.N.G. with support from the German Federal Ministry of Education and Research under Grant Agreements 01GM1922A and 01GM1922B, respectively. A-C.R. under the project ANR-19-RAR4-0001-03. A-C.R. was also supported by the French Fondation Maladies Rares under the project SAM-2022-121409. This work of the Interdisciplinary Thematic Institute IMCBio+, as part of the ITI 2021-2028 program of the University of Strasbourg, CNRS and Inserm, was supported by IdEx Unistra (ANR-10-IDEX-0002), and by SFRI-STRAT’US project (ANR-20-SFRI-0012) and EUR IMCBio (ANR-17-EURE-0023) under the framework of the France 2030 Program. N.D.D received grant support from Deutsche Forschungsgemeischaft (DFG) (DI 2170/3-1 and DI 2170/5-1), Else-Kröner-Fresenius Stiftung (2020_EKES.04) and German Federal Ministry of Education and Research under Grant Agreements 01GM1922A. We acknowledge the Imaging Center of IGBMC, member of the national infrastructure France-BioImaging supported by the French National Research Agency (ANR-10-INBS-04). A-C.R. is a CNRS investigator. N.A. is an Unistra technician personnel. J.N.G. is supported by the HiLF I grant for early career researchers from Hannover Medical School. Some strains were also provided by the CGC, which is funded by NIH Office of Research Infrastructure Programs (P40 OD010440).

We would like to particularly thank the Imaging Center of IGBMC (ici.igbmc.fr) with the assistance of Elvire Guiot, Erwan Grandgirard, Yves Lutz and Bertrand Vernay. We warmly thank Dietmar Manstein (D.J.M) for his significant contributions to the project design and throughout grant application procedures. We also thank Juliette Godin, Michel Labouesse and Patrick Reilly for critical reading of the manuscript. We thank Robert W. Fernandez, post doc in Oliver Hobert’s lab, for training us to NeuroPAL as well as our junior visitor scientists Fanny Bouvier, Emil Zessin and Johann Marois for their help.

## AUTHOR CONTRIBUTIONS

Conceptualisation A-C.R., N.D.D., T.H.; Methodology A-C.R., T.H.; Investigation T.H., N.A., D.S., A.G., G.A., S.Y., F.M., S.Q., J.G., A-C.R.; Writing -Original Draft, A-C.R., T.H.; Writing -Review & Editing, A-C.R., T.H., J.G., N.D.D., S.Q.; Funding Acquisition, A-C.R., N.D.D, D.J.M, J.G.; Resources, A-C.R.; Supervision, A-C.R..

## DECLARATION OF INTERESTS

The authors declare no competing interest.

## SUPPLEMENTAL INFORMATION

Document S1. Figures S1-S6, Tables S1-S3 and supplemental references

Video S1. Cytokinesis arrest, p.R196C embryo. Scale bar = 10μm, related to Figure 3

Video S2. Gastrulation arrest, p.G74S embryo. Scale bar = 10μm, related to Figure 3

Video S3. Ventral enclosure arrest, *rey021* embryo. Scale bar = 10μm, related to Figure 3

Video S4. Extremity burst arrest, p.L65V embryo. Scale bar = 10μm, related to Figure 3

Video S5. Paralyzed arrest, rey021 embryo. Scale bar = 10μm, related to Figure 3

Video S6. Cytokinesis defects, p.R196C embryo. Lifeact::mKate2 (Magenta), NMY-2::GFP (Green). Scale bar = 10μm. Anterior on the left, related to Figure 4

Video S7. Meiosis defects, p.R196C embryo. Lifeact::mKate2 (Magenta), NMY-2::GFP (Green). Scale bar = 10μm. Anterior on the left, related to Figure 4

Video S8. Bleb defects, p.T120I embryo. Lifeact::mKate2 (Magenta), NMY-2::GFP (Green). Scale bar = 10μm. Anterior on the left, related to Figure 4

## STAR★METHODS

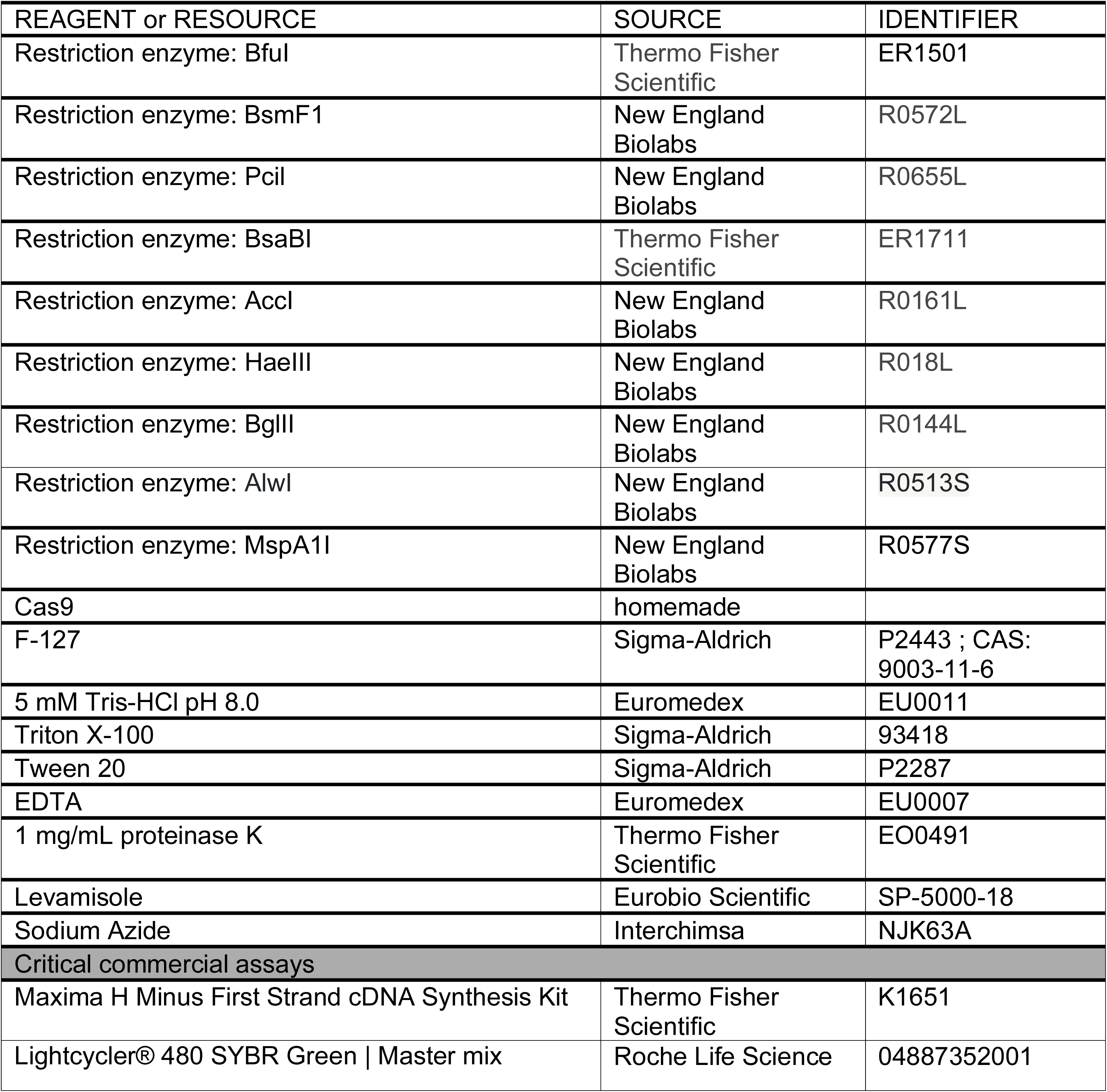
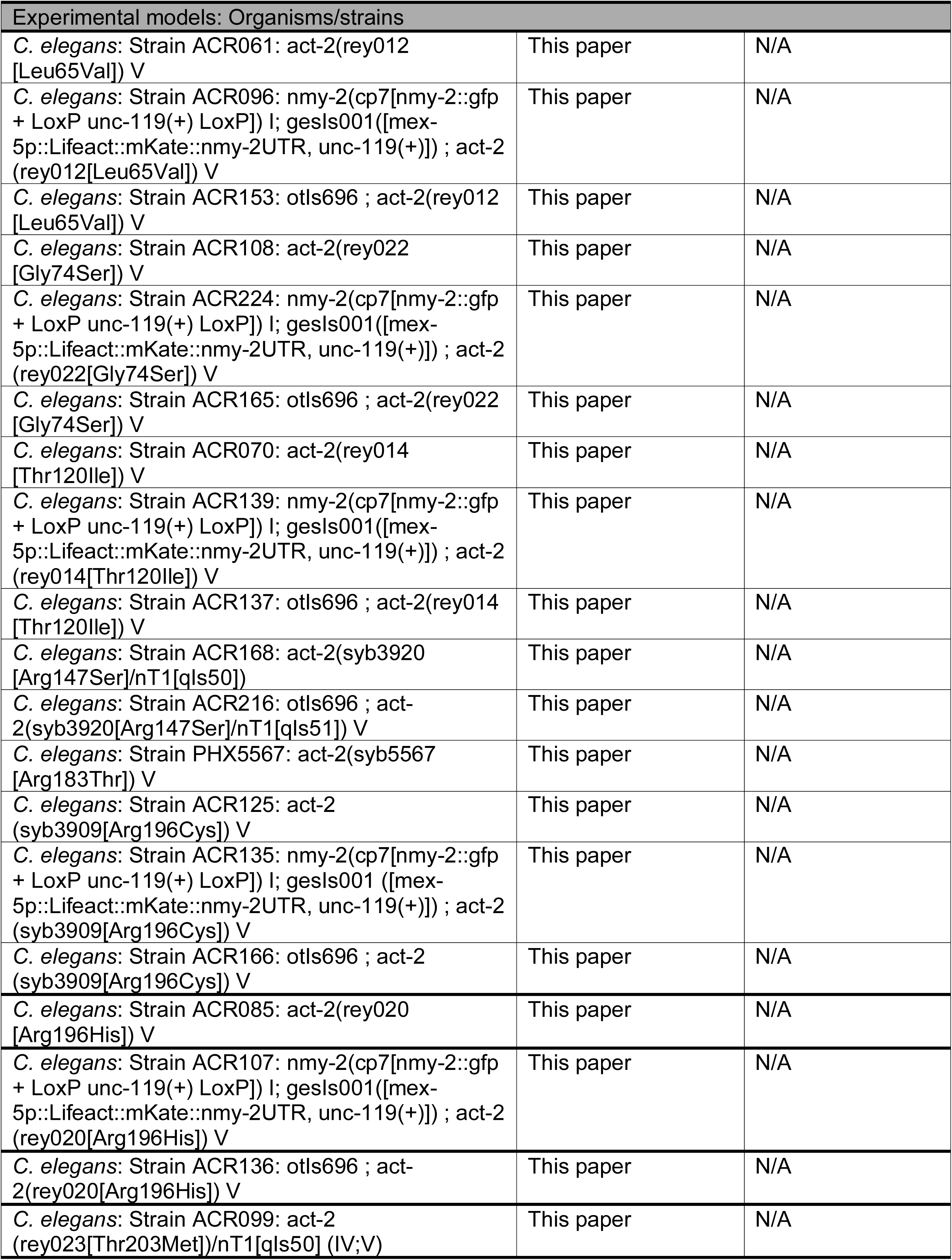
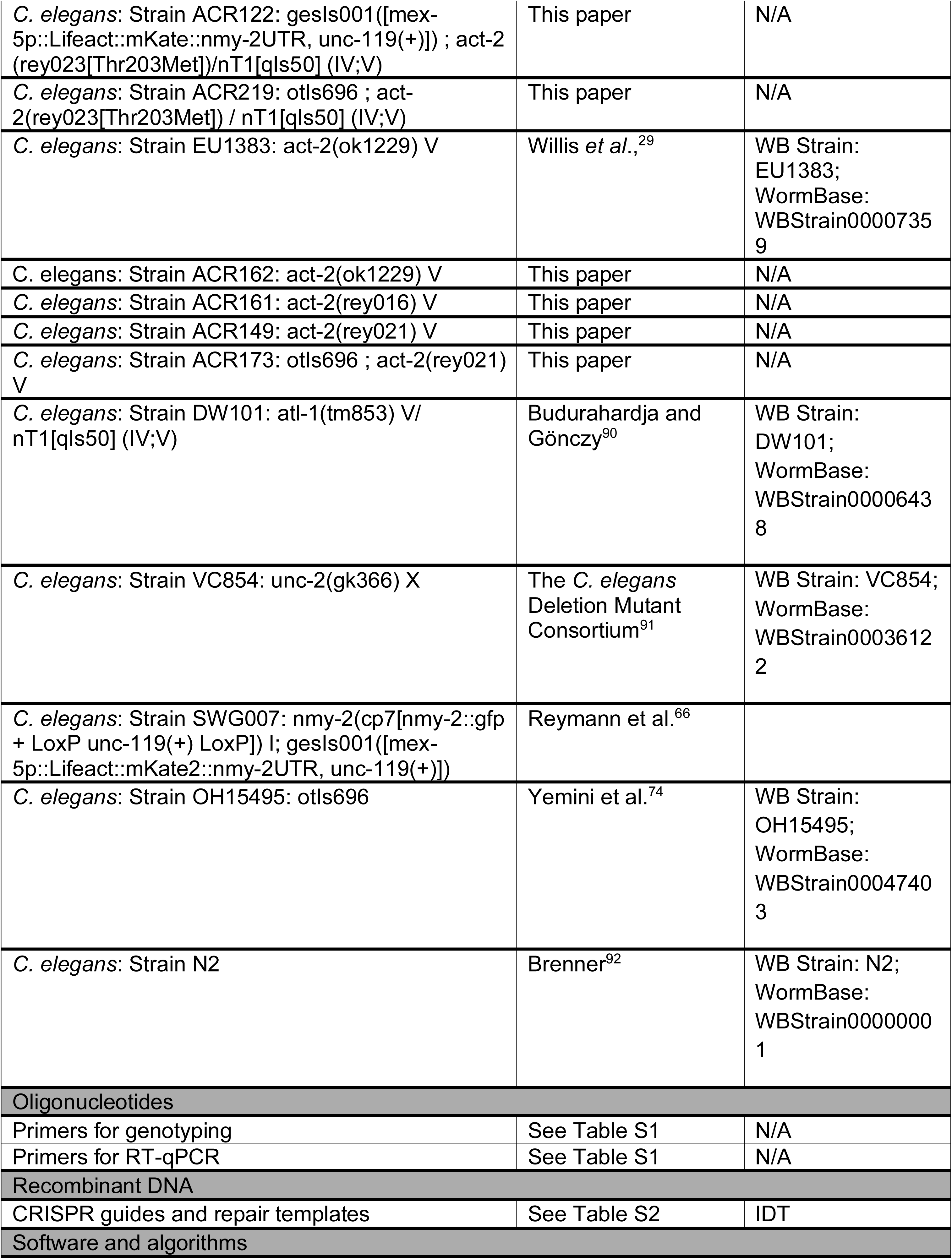
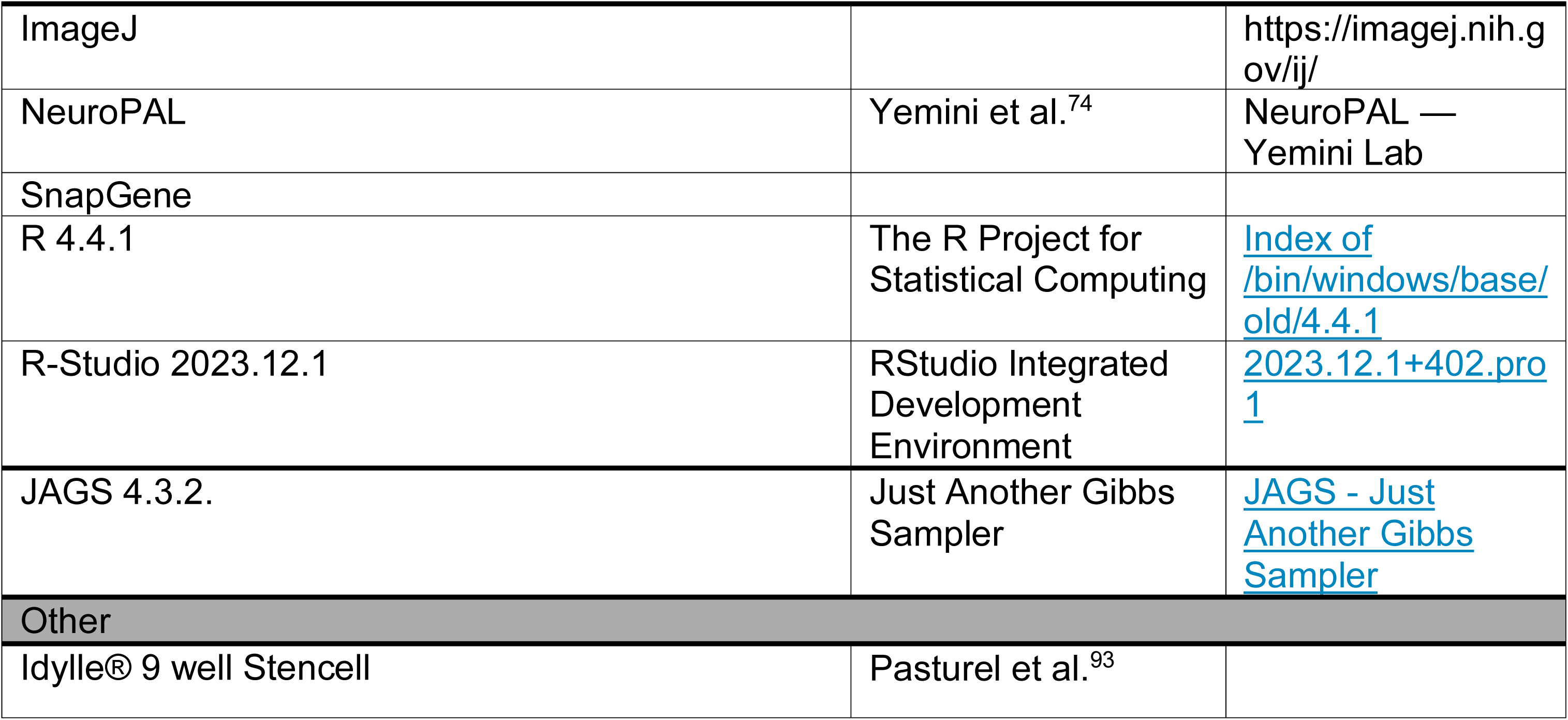
KEY RESOURCES TABLE.

## METHOD DETAILS

### C. elegans handling

Animals were cultured at 20°C on Nematode Growth Medium plates seeded with OP50^92^. Bristol strain (N2) was used as a reference. We thank the Caenorhabditis Genetics Center for providing the strains N2, DW101 and EU1383. All novel actin variants models were generated in the Reymann lab except for two alleles *act-2(syb3920)* and *act-2(syb5567)*, which were outsourced to Suny Biotech®.

### C. elegans generation

Importantly, note that *C. elegans* actins including ACT-2 possesses an additional Cysteine after the initial Methionine in the N-terminus compared to Human ACTB and ACTG1 (ACTB being MDDDIAA and ACT-2 MCDDDVAA, Figure S1A). For simplicity and easier correspondence to patient’s variations, we however referred to *act-2 C. elegans* edited variants using the human amino acid (-1) numbering. For example, human variation ACTB p.L65V is referred to in this publication as *act-2* p.L65V instead of *act-2* L66V in the *C. elegans* frame.

We performed *dpy-10* co-CRISPR editing using purified homemade Cas9 protein, synthetic tracrRNA, and ssODN, as described in Paix et al. 2015^36^. The table of crRNA and repair template used are listed in Supplementary Table 2. Additional silent mutations were introduced either to add or remove a restriction enzyme recognition site for genotyping strategies as well as to mutate PAM or nucleotides within the sgRNA. All generated strains were then outcrossed four times with the N2 strain and all actin coding genes were sequenced to verify that only *act-2* was mutated (genotyping primers available in Supplementary Table 1). Sequencing was performed by Eurofins® and analysed using SnapGene®.

The *act-2(rey016)* allele consists of an insertion-deletion in exon 2 (33bp insertion + 816bp deletion) resulting in a frameshift and a premature STOP codon at the amino acid 70. The *act-2(rey021)* allele presents a 35bp insertion causing a frameshift and a premature STOP codon at the amino acid 120. The *act-2(ok1229)* presents a large deletion starting at the 213th amino acid and extending beyond the 3’ UTR.

### Egg-laying assay

The number of eggs laid by single hermaphrodite animals was quantified on 6-well plates by isolating and transferring individual animals from well to well every 24 hours. Animals were isolated at the L4 stage. After each transfer, eggs laid were manually counted. The number of larvae was manually counted to determine the hatching percentage 36-48 hours after the transfer. This process was repeated over the three days of the time course, otherwise indicated.

Plates were blinded by someone external to the analysis and maintained at the desired temperature (15°C, 20°C, or 25°C) throughout the experiment. Whenever we observed contamination or premature death of the P0 animal, we excluded its dataset from the analysis. Each experiment was performed on around 15 animals including a WT control group on the same medium pour.

Regarding figure 2D and S3A/C, we generated heterozygote animals by crossing L4-staged homozygous actin variants animals with N2 males and quantified the brood size of F1 animals (2E and S3B/D). After experiment completion, we verified each animal’s genotype by sequencing their progeny (genotyping primers are available in Supplementary Table 1). In figure S3C/D heterozygous animals were obtained after a cross between N2 males and p.R147S/nT1 and p.T203M/nT1 heterozygous animals; but only non-nT1 (ie non-GFP fluorescent pharynx) animals were considered —hence the R147S/+ and T203M/+ maternal genotype.

### 3D motility assay

The ability of animals to move in a 3D medium was assessed using a pluronic-based burrowing assay as described in Lesanpezeshki et al ^73^ and optimized by Ana Carvahlo (personal communication) as follows. Liquid pluronic gel was made by adding 13g of F-127 powder into 37mL of Ultra Pure water and leaving the solution at 4°C for 48 hours. At least an hour before the experiment, pluronic gel and consumables were put at 10°C and kept refrigerated unless otherwise stated. Animals were isolated at the L4 stage 24 hours before the experiment and blinded by someone external to the experiment. Each experiment included a WT control group.

For each experiment, we put a 25µL drop of liquid pluronic gel in each well of a 24 -well plate and added 10 animals into each drop. We left the plate at 20°C for a few minutes for the drop to set. Next, we added 1.3mL of liquid pluronic gel to each well to obtain a gel height of 0.6µm. After waiting 10 minutes for the gel to solidify, we added a 10µL drop of fresh concentrated (100mg/mL) OP50 to use as a chemo-attractant and kept the plate at 20°C. Then, the number of animals that reached the well’s top was manually counted every 10 minutes for two hours.

The negative control, *unc-2(gk366)* is a knockout of the Human CACNA1B (calcium voltage-gated channel subunit alpha1 B) ortholog, thus predicted to enable high voltage-gated calcium channel activity expressed in whole body wall muscle and required for proper animal motility^94^.

### Chemo-sensation assay

To ensure that the 3D motility defects observed were unrelated to the animals’ potential inability to sense food, we performed a standard chemosensation assay. Animals, isolated at the L4 stage 24 hours before the experiment were blinded by someone external to the experiment. Animals were washed 4 times in M9 and placed at the center of a 6cm plate NGM plate without food. Fresh concentrated (100mg/mL) OP50 and Ultra Pure H_2_O were placed on opposite extremities of the plate. After one hour, the number of animals on each side of the plates was counted as well as the animals that did not move from their original positions (“Immobile animals” in Figure S5). The chemotaxis index was calculated using 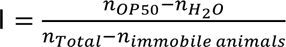.

### RT-qPCR

Single animal RT-qPCR was performed as in Ly, Reid and Snell ^55^. An hour before the experiment, we isolated L4 animals on a plate without food. Single L4 animals were transferred in 0.2mL Eppendorf tubes containing 2µL of lysis buffer (5 mM Tris-HCl pH 8.0, 0.5% Triton X-100, 0.5%, Tween 20, 0.25 mM EDTA and 1 mg/mL proteinase K) without sticking them with bacteria. Tubes were incubated at 65°C for 10 min, then at 85°C for 1min in a T100TM Thermal Cycler. After lysis completion, we used the Maxima H Minus cDNA synthesis kit. Genomic DNA was removed by adding 1µL of a master mix containing 1µL of dsDNAse buffer, 1µL of DsDNAse, and 8µL of nuclease-free water. The solution was incubated at 37°C for 2 minutes and immediately chilled on ice. cDNA synthesis was generated by adding 1µL of a mix containing 0.5µL oligo dT primer, 0.5µL 10mM dNTP, 6.5µL nuclease-free water, 2µL RT buffer, and 0.5µL Maxima H Minus enzyme mix to the lysate. Tubes were centrifugated and incubated them at 25°C for 10 minutes, then at 55°C for 30 minutes, and finally 5 minutes at 85°C. We diluted the cDNA to 25 µL with nuclease-free water and used it immediately. We prepared 6 mixes of primer (*act-1, act-2, act-3, act-4, act-5,* and *iscu-1*) containing 30.6µL of PCR grade water, 7.2µL of the forward primer at 10µM, 7.2µL of the reverse primer at 10µM (Available in Supplementary Table 1) and 90µL of the Lightcycler® 480 SYBR Green | Master mix. We added 7.5µL of primer mix for 2.5µL of lysis into a 384-well plate. We sealed the plate using the provided foil with gloves and centrifuged it at 1500xg for 2 minutes. We used a LightCycler® for qPCR following these steps; firstly a pre-incubation at 95°C for 15 minutes with a 4.4 ramp, secondly a step of amplification of 45 cycles composed of 10 seconds at 95°C with a 4.4 ramp, 15 seconds at 54°C with a 2.2 ramp and 10 seconds at 72°C with a 4.4 ramp. Thirdly, a melting curve analysis composed of 5 seconds at 95°C with a 4.4 ramp, 65 seconds at 70°C with a 2.2 ramp, and continuous heating to reach 95°C before cooling back at 40°C.

The Cts measured by the Lightcycler® were exported to a text file and were analysed using R and Rstudio (see Key resources table). Values were normalised using the Ct values of the reference gene *iscu-1* ^55,95^ and to the N2 results of the same experiment. Some animals were excluded from analysis when their Ct values reached 40. Primer pairs efficiencies were measured using standard curve analysis available in Figure S2.

### Microscopy

#### Embryonic development time-lapse imaging

L4 animals were put on a plate at 20°C 24h before acquisition. To obtain embryos, dissections were performed on a slide covered with an Idylle® 9 well Stencell^93^ with 5µL of M9 and 4 animals of the desired genotype in each well. Then, corpses were removed from each well using a mouth micropipette. The slide containing the dissected animals was then stuck to a #1.5 coverslip (Marienfeld Superior) and placed on a Zeiss Axio-Observer 7. DIC images were then acquired using a Hamamatsu orca flash 4.0 LT camera, every 10 seconds for 40 minutes and then every 10 minutes for at least 14 hours with a 20x/0.8 PLAN-Apochromat objective. Given the low frequency of some of the observed phenotypes, we deliberately chose to favour high throughput with low magnification at the expense of image quality: 9 wells were simultaneously acquired, thereby allowing replicates of 3 mutant conditions and 3 WT controls, representing in total ∼20 to 25 (x,y) positions containing each 2 to 15 embryos simultaneously observed during overnight acquisitions. The room temperature was controlled and set to 20-22 degrees.

### Early embryogenesis

L4 animals were put on a plate at 20°C 24 hours before acquisition. Dissection of 5 animals was performed in 5µL of M9. The slide containing the dissected animals was then placed on a 2% agarose pad and placed on an inverted microscope Nikon EclipseTi-2 equipped with YOKOGAWA CSU X1 spinning disk head. Pictures were then acquired using a Photometrics Prime 95B camera each 2 or 10 second for 15 minutes with a plan apo 100x/1.45 Nikon objective.

### Gonad imaging

L4 animals were immobilised in 10 µL of 1mM Levamisole and placed on a 2% agarose pad. Acquisition of gonads was performed on a LEICA DMI8 inverted microscope equipped with Yokogawa CSU W1 Spinning disk. Z-stacks were acquired using a Hamamatsu Orca Flash 4.0 camera with a 40X HCX PL APO oil immersion objective with a 0.2µm Z-step. Gonad widths were measured using FIJI by drawing a line with a 120-pixel width across the gonads germs cells and rachis in front of the +1 oocyte. Classification of phenotypes was performed visually as represented in ^58^. Heatmap was built using the heatmap.2 function from the R package gplots. The mean number of observed phenotypes for each variable was scaled in the row direction.

### NeuroPal

L4 animals were immobilised in 7µL of 125mM sodium azide and placed on a 2% agarose pad. Acquisitions were made accordingly to https://www.hobertlab.org/wp-content/uploads/2019/09/Configuring-Your-Microscope-for-NeuroPAL-v2.pdf on a Confocal Inverted Leica TCS SP8X equipped with DMI6000 microscope White Light Laser using a 40x/1.3 HC PL APO CS oil immersion objective with a distance of 0.9µm between each Z. Mapping and comparison of the expected expression pattern is performed using the NeuroPAL software and reference atlas map.

### Principal Component Analysis

Principal Component Analysis (PCA) was computed and plotted using the FactoMineR package with the PCA function using 9 dimensions, we set the ‘graph’ argument to true to obtain the variables factor map presented in Figure S6A. Missing values were replaced by the mean of the corresponding variable. Matrices show the quality of representation of variables across dimensions and were constructed using the get_pca_var function from the FactoMineR package.

We exported the coordinates of each point and calculated the distance between each point and the WT in the two first dimensions.

### *In silico* analysis

The primary sequence of ACT-2 from *C. elegans* was obtained from the UniProt database (ID: P10984). We used Alphafold2 ^96^ to predict the structure of ACT-2 G-actin. Calculations were performed on the Halime HPC cluster at the Institute for Biophysical Chemistry at Hannover Medical School. We visualized ATP in the nucleotide binding cleft of ACT-2 G-actin by aligning the predicted ACT-2 structure with the structure of bovine ATP-bound G-actin (PDB-ID: 2BTF, ^97^). The homology model of the ACT-2 filament was generated using the SWISS-MODEL server ^98^ with the ADP-bound human cytoskeletal β-actin filament (PDB-ID: 8DNH, ^99^ as a template. ChimeraX ^100^ was used for visualization of the structures and assessment of the potential implications of the variants for actin structure.

## QUANTIFICATION AND STATISTICAL ANALYSIS

We used R 4.4.1 through R-Studio 2023.12.1 and JAGS for all data analysis with all the required additional packages in their latest version available. All original code has been deposited and is publicly available via our Gitlab (https://gitlab.com/igbmc/reymannlab/2025-hecquet-iscience). The statistical analyses were performed in the frame of Bayesian and frequentist methods.

### Bayesian Framework

The experimental data was enriched within the Bayesian framework by inference based on posterior distribution description. For embryonic lethality: Poisson distribution was used to estimate the embryonic lethality rates using an offset to account for the dependency with the number of eggs laid per animal (brood size) and specific parameters values for each genotype. Parameters were either given informative Normal priors: considering a very small lethality rate for WT N(0.01, 0.1), more probable lethality for all mutants N(0.1, 0.1) except the KO homozygotes for which we estimated a possible lethality of half the embryos N(0.5, 0.1). The analysis was also computed using low informative priors for all genotypes N(0,0.01) and gave similar results within a thousandth. Note that the observed 100% embryonic lethal genotypes were not included in the model. The posterior distribution was computed using the Markov chain Monte Carlo method (McMC). A burn-in of 2000 iterations, followed by 50000 iterations was used for each of the 3 McMC chains, yielding a final 28800 iterations-sample for retrieving posterior distribution characteristics. Convergence of the McMC sample chains was checked graphically. Convergence was observed in each case. For each analysis on the main outcome, we computed the 95% credible interval of the embryonic failure rates for each genotype.

Concerning the 3D motility assay, we compared the observed success rate to reach the chemoattractant after 2h by calculating their respective posterior distributions using two binomial distributions. We used the following priors: for WT mean = 0.9, sd=0.1 thus a probability modelled as p∼dbeta(7.2, 0.8) and for all mutants a mean=0.5, sd=0.2 thus a probability modelled as p∼dbeta(2.65, 2.65). A burn-in of 5000 iterations, followed by 105000 iterations was used for the McMC chain, in order to retrieve the posterior distributions. The odds ratio (OR), an estimation of the multiplication of the risk to fail reaching the target compared to WT, as well as the probability (p) that the difference between the two mean posterior probabilities is superior to 0.2 are indicated.

### Frequentist Analysis

We used the R package rstatix for all frequentist statistical analyses. For multiple mean comparisons, we first verified the equality of variances and normality distribution of groups using a Levene and Shapiro test respectively. If those were verified, we realized an ANOVA test followed by a Tukey’s range test. Otherwise, we realized a Kruskall-Wallis followed by a Dunn test. P-values obtained with the Dunn test were adjusted using the Bonferroni method.

Statistics of Figure 5A were done on the values at the 2-hour timepoint. A summary of all tests and p-values is available in Supplementary Table 3.

---

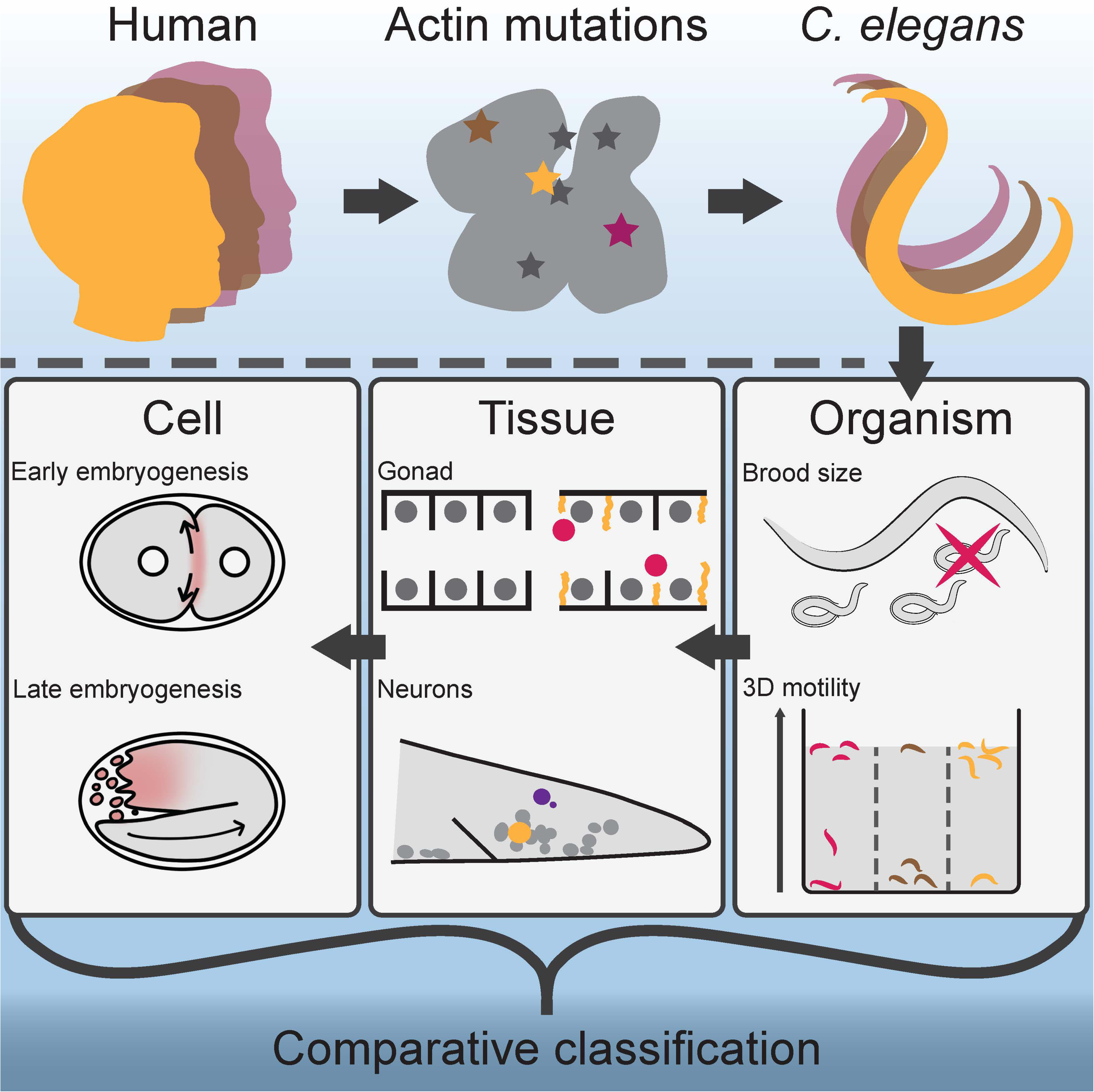

**Figure S1.**
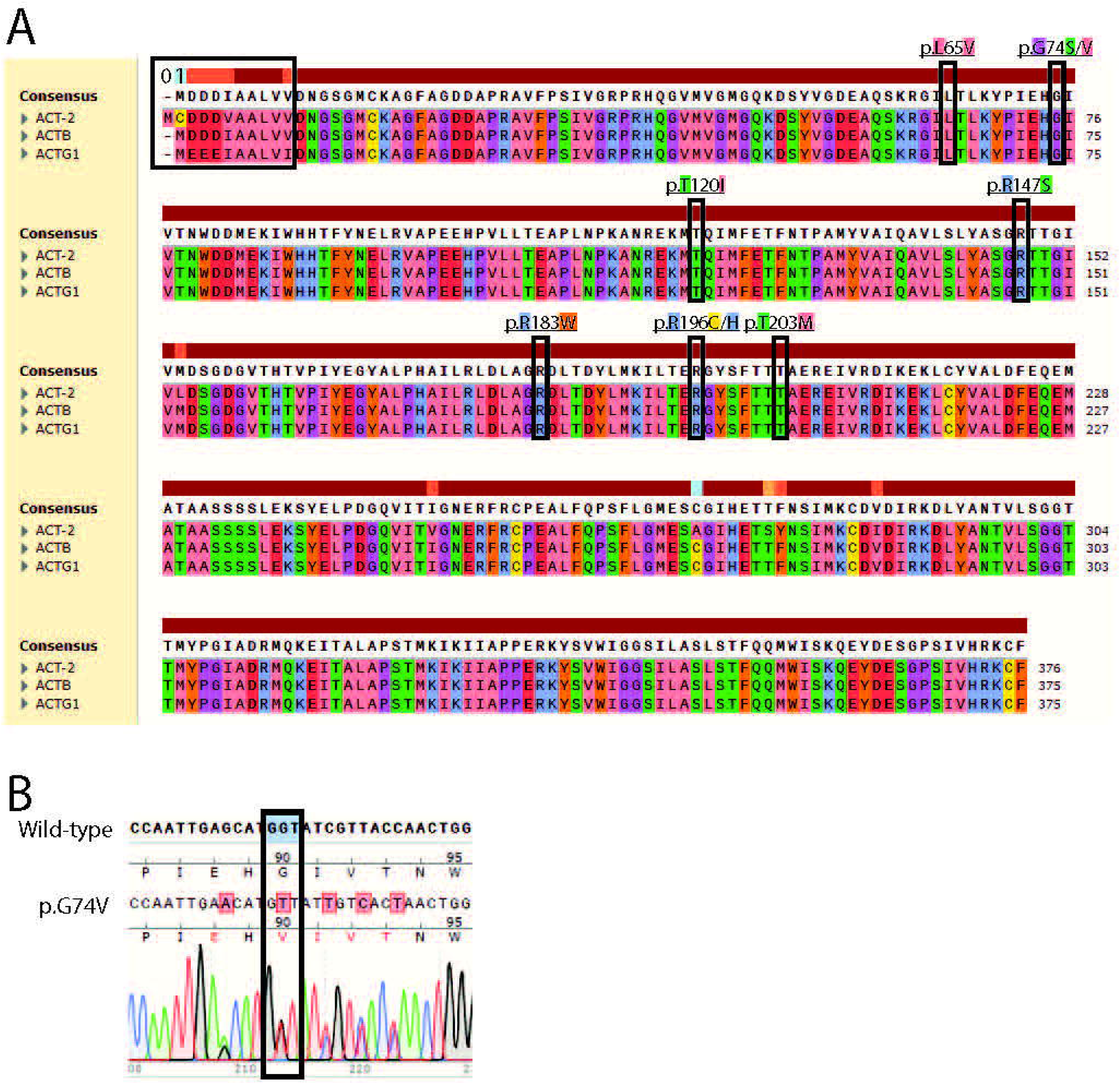

**Figure S2.**
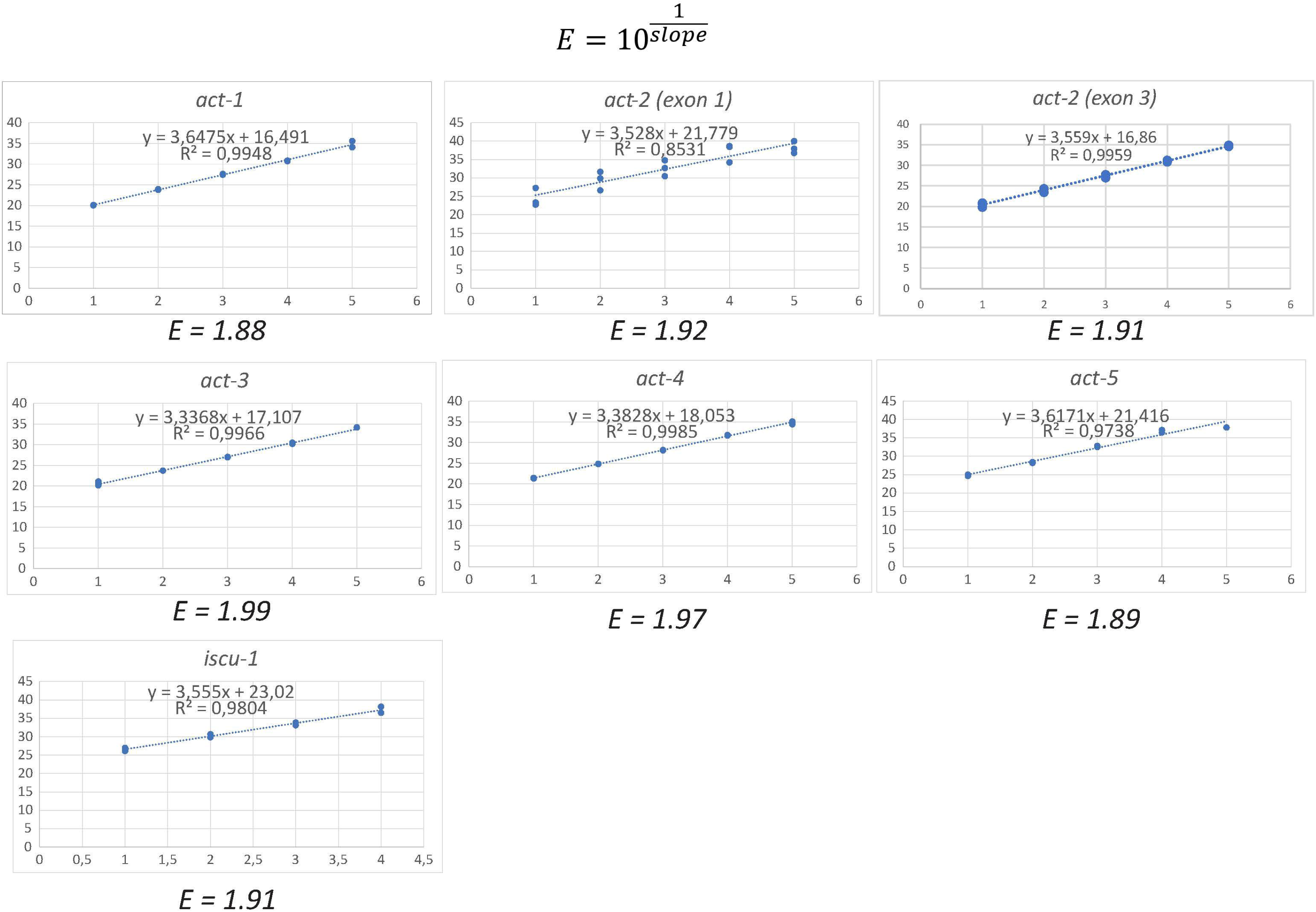

**Figure S3.**
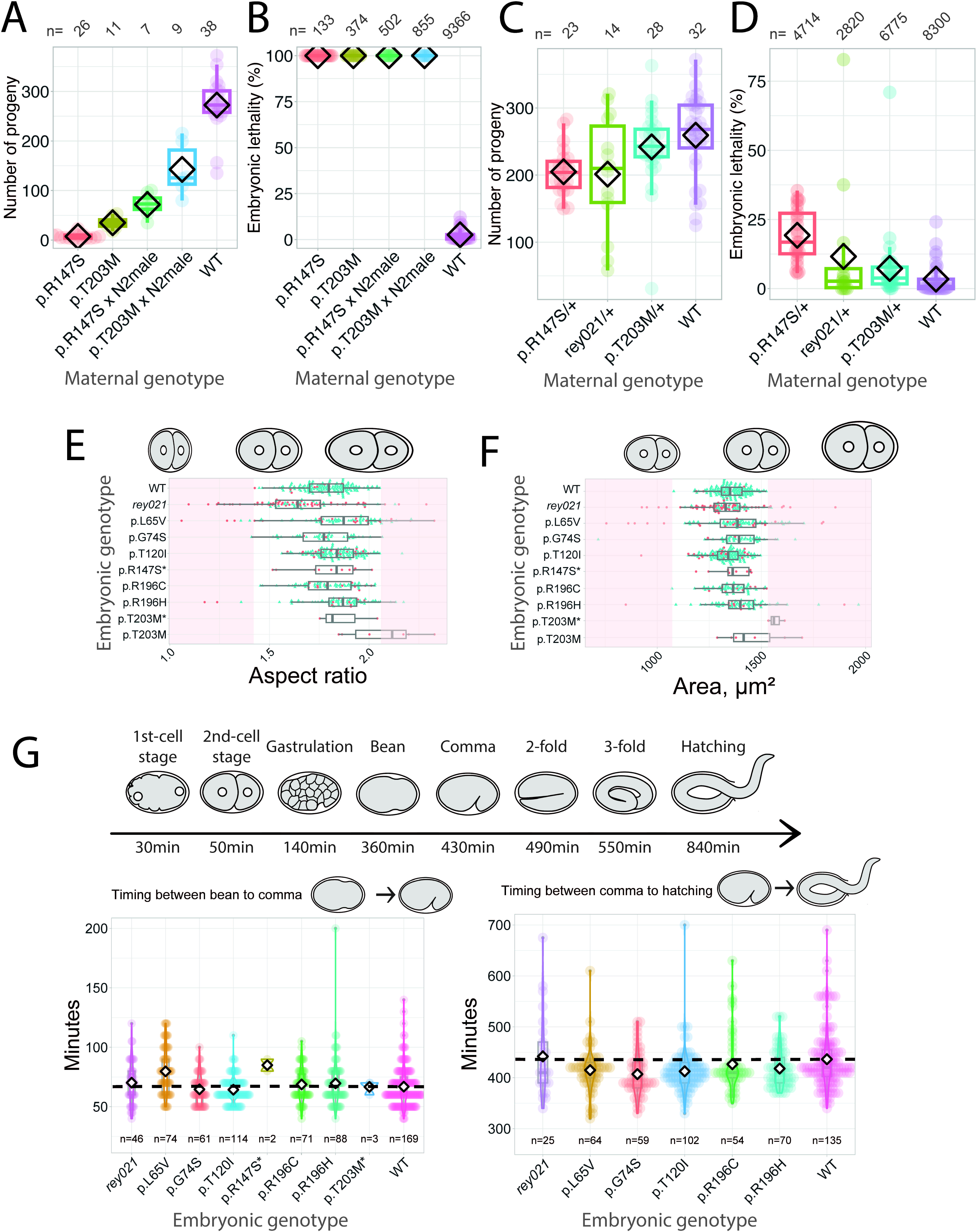

**Figure S4.**
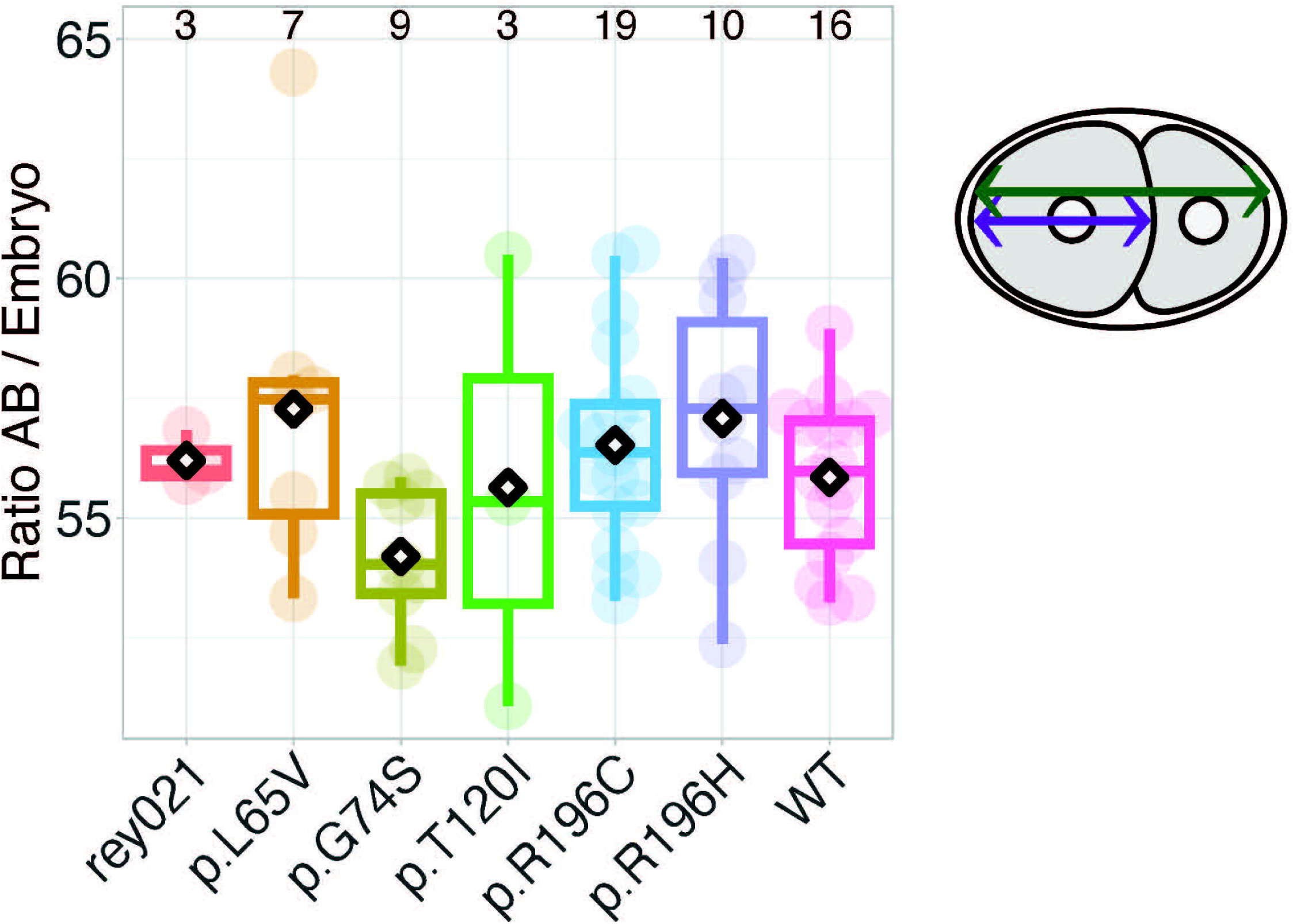

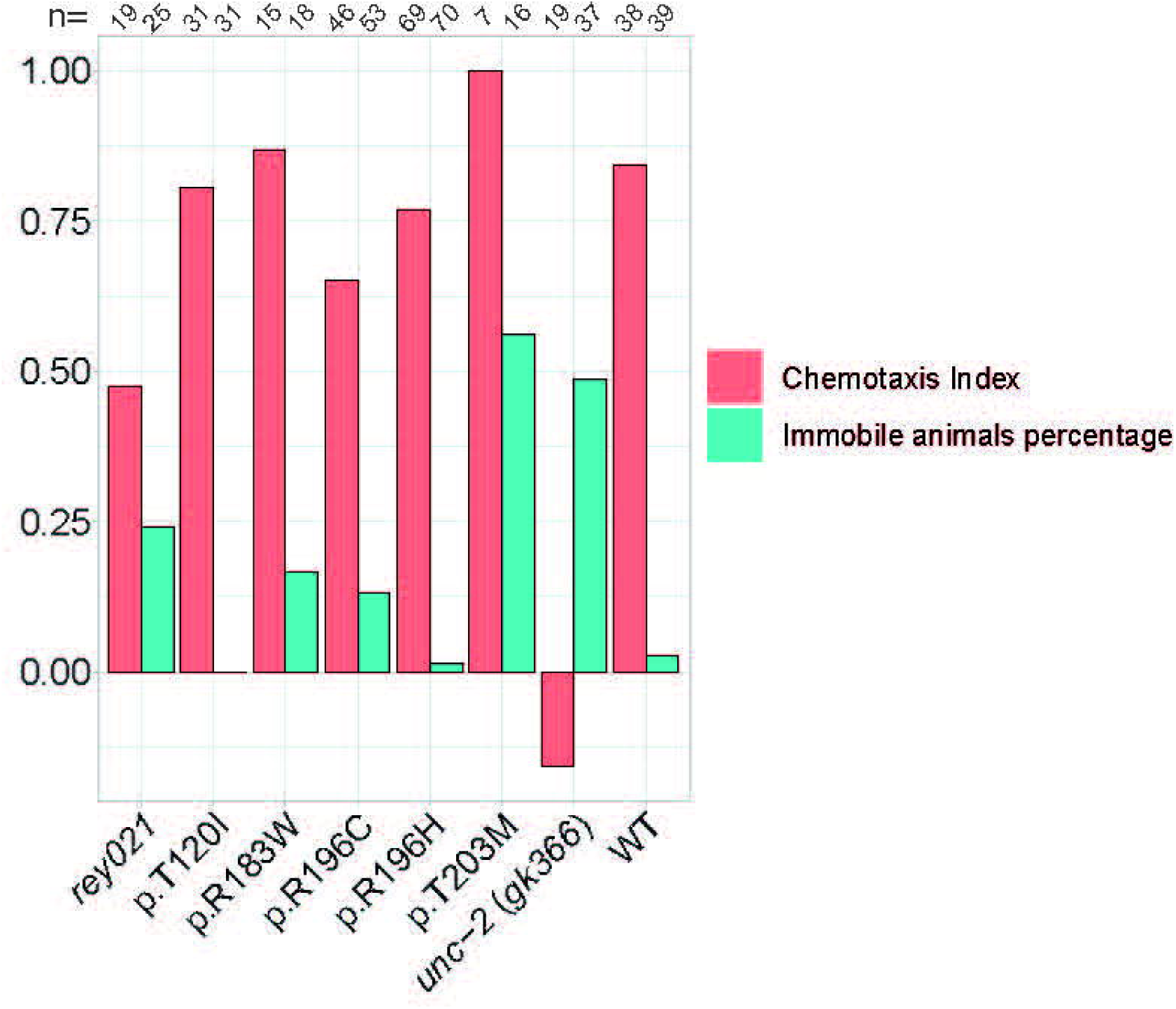
aaaa

**Figure S6.**
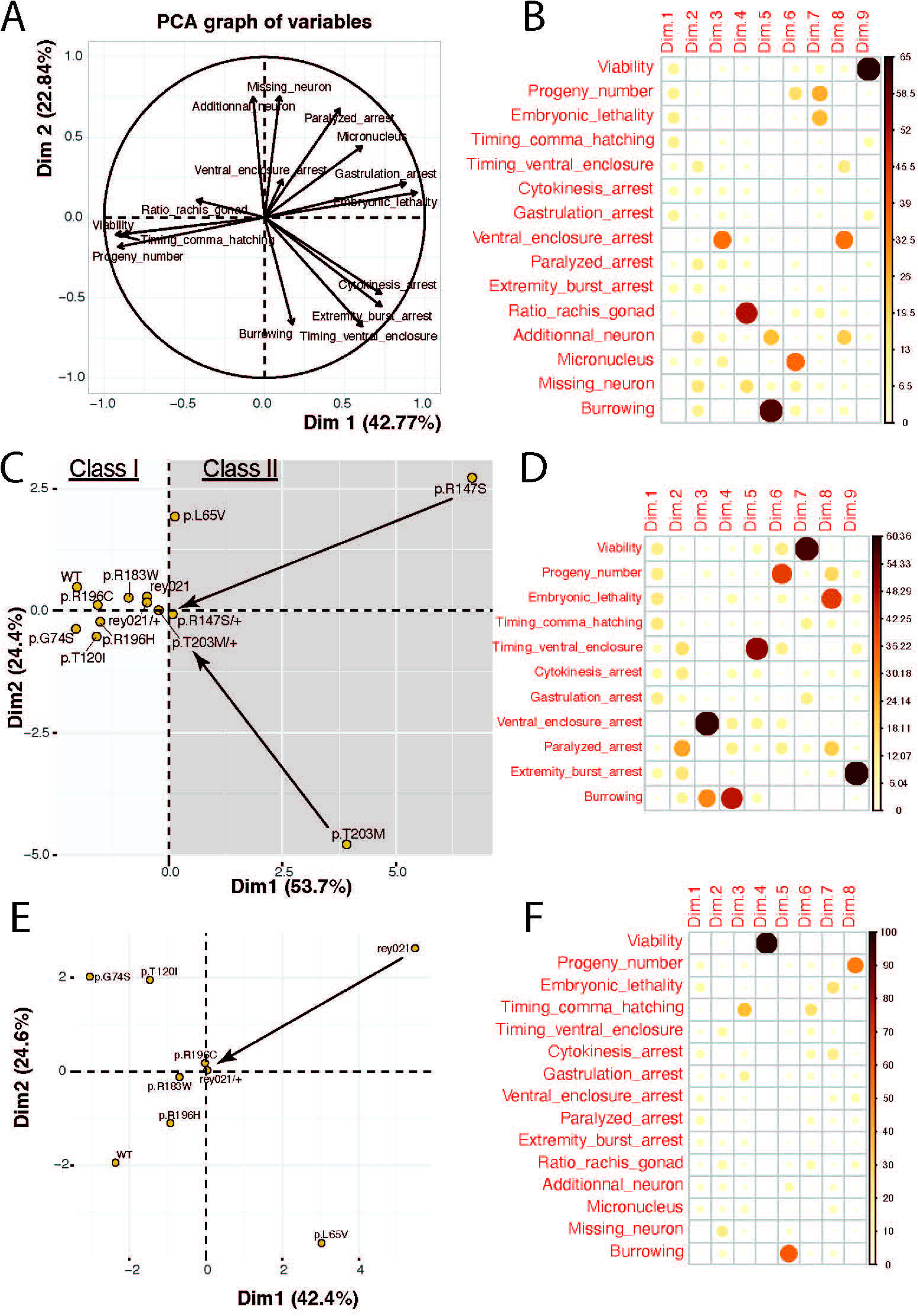

## Notes

### Competing Interest Statement

The authors have declared no competing interest.

### Summary of Updates

author affiliation updated ; Figure 2 and Figure S3 were revised and some sections of the result and discussion were updated accordingly ; minor correction for typos.

