## Supplement Files for "Multiscale characterization of *Caenorhabditis elegans* mutants to probe functional mechanisms of human actin pathological variants"

**Figure S1. Similarity of human and *C. elegans* actins, Related to Figure 1**

(A) Amino acid alignment of *C. elegans'* ACT-2, human ACTB and ACTG1 showing high conservation of amino acid as well as amino acid physico-chemical properties (Snapgene). Variants positions are framed with black boxes. (B) Alignment of sequences obtained from the sequencing of F1 animals from the CRISPR/Cas9 *act-2* p.G74V generation show the presence of the Gly to Ser variant (Snapgene, black box). Top rows show the reference *act-2* sequence and its translation. Bottom rows show the obtained sequence, peaks and predicted translation.

**Figure S2. Efficiencies of primers used in the RT-qPCR experiments, Related to Figure 2**

(A) Standard curve analysis to determine the efficiency of amplification of the 6 primer pairs used for RT-qPCR. Efficiency is calculated using $E = 10^{\frac{1}{slope}}$.

**Figure S3. Heterozygous *act-2* animal brood sizes and eggshell measurements, Related to Figure 3**

(A) Total progeny number counted per hermaphrodite for indicated maternal genotypes: homozygous p.R147S and p.T203M variants after mating with N2 males. (B) Percentage of eggs that failed to hatch from these mated actin variant hermaphrodites. The percent of dead eggs for one animal is shown as a dot, overlaid by standardised boxplots and mean values are represented by a diamond. (C) Total progeny number were counted per heterozygous *act-2* variant animals. The sum of eggs laid for one animal is shown as a dot, overlaid by standardised boxplots and mean values represented by a diamond. (D) Percentage of eggs that failed to hatch. The percent of dead eggs for one animal is shown as a dot, overlaid by standardised boxplots and mean values are represented by a diamond. (E, F) Red boxes show the area where no WT eggshells are present. Red points show embryos that did not complete embryogenesis while blue triangles represent a successful embryogenesis. Embryo schemes above are at the right scale. (E) Aspect ratio measurement of embryos' eggshell. Aspect ratio is defined as the ratio between the major and minor axis. (F) 2D Area measurement of embryos' eggshell. E. Timeline of recognizable morphogenesis stages during *C. elegans'* embryogenesis. Time counted after fertilization, from Hall and colleagues [S1]. Leftmost plot presents the timing in minutes between the bean stage and the comma stage. Rightmost plot presents the timing in minutes between the comma stage and hatching. * embryos coming from heterozygous balanced animals.

**Figure S4. Cell sizes at the embryonic two-cell stage, Related to Figure 4**

Size of the anterior cell AB (Magenta arrow) divided by the embryo's size (Green arrow) in percent. Boxplots display: the maximum and minimum values, the median as well as first and third quartiles.

**Figure S5. Chemotaxis assay, Related to Figure 5**

Chemotaxis assay. The chemotaxis index is calculated $I = \frac{n_{OP50} - n_{H_2O}}{n_{Total} - n_{immobile\ animals}}$ and show the proportion of animals chemoattracted to concentrated OP50 (100mg/mL) (in red). Blue bars show the percentage of animals that did not move from where they were placed for the assay.

**Figure S6. Variables used for the PCA, Related to Figure 6**

(A) Variables factor map used to build the Principal Component Analysis plot in the two first dimensions. The Variables factor map presents a view of the projection of the observed variables projected into the plane spanned by the first two principal components. (B) Matrix of variables presented in Figure 6A, as well as their quality of representation in each dimension (cos2). (B) Principal Component Analysis plot of analysed actin variant animals using only the variables quantified without introducing fluorescent reporters in the actin variant background. Arrows link homozygotes to their respective heterozygotes. (D) Variables presented in S6C, as well as their quality of representation in each dimension (cos2). (E) Principal Component Analysis plot of analysed actin variant animals without p.T203M and p.R147S variations. (F) Variables presented in S6E, as well as their quality of representation in each dimension (cos2).

**Table S1. Primer sequences.**

| *Gene* | Forward | Reverse | Hybdridization temperature | Amplicon length (pb) | Purpose |
|---|---|---|---|---|---|
| *act-1* | ttccccacgtgttctgtg | GTTTGTGCAAGTTGACGAAG | 54°C | 1323 | Genotyping |
| *act-2* | CTCGTAGTTGACAATGGATC | CCAACAGTGATGACTTGTCCG | 56°C | 776 | Genotyping |
| *act-2* | gctggtttaattggaaagct | gtcagccagttcaccacgaa | 55°C | 2544 | Genotyping |
| *act-3* | ccttcccacacccacttg | gggaccataacgatttggtg | 60°C | 1495 | Genotyping |
| *act-4* | ctcgaccgaattctcgcc | ggttgcgggataatacgaac | 50°C | 1379 | Genotyping |
| *act-5* | CGTTTTCTATAAACTCGACTTC | GAAATGGAAGGCACATGT | 51°C | 1500 | Genotyping |
| *act-1* | GCCCCATCAACCATGAAG | gtttgtgcaagttgacgaag | 54°C | 196 | RT-qPCR |
| *act-2(exon3)* | GCCCCATCAACCATGAAG | gacgttatctggacatttcc | 54°C | 226 | RT-qPCR |
| *act-2(exon1)* | CTCGTAGTTGACAATGGATC | CCTCTCTTGGATTGGGCC | 54°C | 167 | RT-qPCR |
| *act-3* | GCCCCATCAACCATGAAG | ggtaaggcgaagagcTTAGA | 54°C | 183 | RT-qPCR |
| *act-4* | ATCGATTGTCCACCGCAAG | ggttgcgggataatacgaac | 54°C | 84 | RT-qPCR |
| *act-5* | GCTCCAAGCACAATGAAG | gaaatggaaggcacatgt | 54°C | 215 | RT-qPCR |
| *iscu-1* | GAAGGTTATCGATCACTACGAG | CGTTATCGTCGACTCGAATC | 54°C | 137 | RT-qPCR |

**Table S2 List of guides and repair template sequences.**

Lower case (non-coding), upper case (coding), red (silent mutation), blue (non-synonymous mutation of interest), bold (PAM sequence), underlined (crRNA sequence).

| *Mutation and allele name* | crRNA | Repair | Enzyme used for Genotyping |
|---|---|---|---|
| *p.L65V, rey012* | CAAUUGGGUAC UUAAGGGUA | Ggtaaattttcaaaaaattt g acc gat tgg aaa tag ttg ttt tta ggt **ATC** **G**TC ACT CTT AAA TAC CCA ATT GAG CAT GGT ATC G | BfuI |
| *p.G74S, rey022* | CGUUACCAACU GGGACGACA | CTT ACC CTT AAG TAC CCA ATT GAG CAT AGT ATC GTG ACA AAT TGG GAT GAC A**TG G**AA AAA ATC TGG CAT CAC | BsmF1 |
| *p.G74V* | AGCAUGGUAUC GUUACCAAC | g ttt tta gGT ATC CTT ACC CTT AAG TAC CCA ATT GAA CAT GTT ATT GTC ACT AAC **TGG** GAC GAC ATG GAA AAA ATC TGG C | PciI |
| *p.T120I, rey014* | AUACAUGGCUG GGGUAUUGA | CT AAC CGT GAA AAG ATG ATC CAA ATC ATG TTC GAG **ACA** TTC AAT ACA CCA GCA ATG TAC GTC GCC ATC CAA GCT G | BsaBI |
| *p.R147S, syb3920* | N/A | CAA GCT GTC CTC TCC CTC TAC GCT TCC GGG TCG ACC ACG GGA ATC GTC CTC GAC TCT GGA GAT | AccI |
| *p.R183W,*<br><br>*syb5567* | N/A | ACG CAC ACA GTC CCA ATC TAC GAA GGA TAT GCC CTC CCA CAC GCC ATC CTC CGT CTT GAC TTG GCC GGA TGG GAT CTT ACT GAT TAC CTC ATG AAG ATC CTT ACC GAG CGT GGT TAC TCT TTC ACC ACC ACC GCT GAG CGT GAA ATC GTC CGT GAC ATC AAG GAG AAG CTT TGT TAC GTC GCG | HaeIII |
| *p.R196C, syb3909* | N/A | GGA ATC GTC CTC GAC TCT GGA GAT GGT GTT ACG CAC ACA GTC CCA ATC TAC GAA GGA TAT GCC CTC CCA CAC GCC ATC CTC CGT CTT GAC TTG GCT GGA CGT GAT CTT ACT GAT TAC CTC ATG AAG ATC TTG ACC GAA TGC GGT TAC TCT TTC ACC ACC ACC GCT GAG CGT GAA ATC GTC CGT GAC ATC AAG GAG AAG CTT TGT TAC GTC GCG CTC | BglII |

| | | | |
|---|---|---|---|
| | | GAT TTC GAG CAA GAA ATG GCC ACC GCC | |
| *p.R196H, rey020* | AUGAAGAUCCU UACCGAGCG | CTT ACT GAT TAC CTC ATG AAA ATT CTT ACC GAG CA**T GG**T TAC TCT TTC ACC ACC ACC GCT GAG CGT GAA ATC | AlWI |
| *p.T203M, rey023* | GACGAUUUCAC GCUCAGCGG | G AAG ATC CTT ACC GAG CGT GGT TAC TCT TTC ACC **ACT** ATG GCA GAG CGA GAA ATT GTG CGT GAC ATC AAG GAG | MspAII |
| *KO, rey021* | AGCAUGGUAUC GUUACCAAC | N/A | |
| *KO, rey016* | AUACAUGGCUG GGGUAUUGA | N/A | |

**Table S3 : Frequentist statistics summary.**

| Related to Figure 2D, Kruskall-Wallis and Dunn test with adjusted p-value. | | | | | |
|---|---|---|---|---|---|
| | WT | *rey021/rey021* | *rey021/+* | *rey016/rey016* | *rey016/+* |
| WT | – | 1.3e-6 | 1 | 2.3e-5 | 1.54e-2 |
| *rey021/rey021* | – | – | 9.4e-2 | 1 | – |

| Related to Figure 2E, Kruskall-Wallis and Dunn test with adjusted p-value and extreme outliers. | | | | | |
|---|---|---|---|---|---|
| | WT | *rey021/rey021* | *rey021/+* | *rey016/rey016* | *rey016/+* |
| WT | – | 4.37e-12 | 1 | 3.50e-3 | 1 |
| *rey021/rey021* | – | – | 7.24e-5 | 1.58e-1 | – |

| Related to Figure 2E, Kruskall-Wallis and Dunn test with adjusted p-value and no extreme outliers. | | | | | |
|---|---|---|---|---|---|
| | WT | *rey021/rey021* | *rey021/+* | *rey016/rey016* | *rey016/+* |
| WT | – | 3.97e-12 | 1 | 2.82e-3 | 1 |
| *rey021/rey021* | – | – | 3.47e-6 | 1.77e-1 | – |

| Related to Figure 3A, Kruskall-Wallis and Dunn test with adjusted p-value. | | |
|---|---|---|
| | WT | *rey021* |
| *rey021* | 4.29e-6 | – |
| p.L65V | 1.02e-1 | 1 |
| p.G74S | 1 | 3.99e-2 |
| p.T120I | 1.84e-2 | 1 |
| p.R147S | 1.62e-10 | 1 |
| p.R183W | 1 | 3.56e-2 |
| p.R196C | 1 | 4.23e-1 |
| p.R196H | 1 | 4.01e-1 |
| p.T203M | 2.39e-7 | 1 |

| Related to Figure 3B, Kruskall-Wallis and Dunn test with adjusted p-value and extreme outliers. | | |
|---|---|---|
| | WT | *rey021* |

| | | |
|---|---|---|
| *rey021* | 1.49e-5 | – |
| p.L65V | 1.07e-2 | 1 |
| p.G74S | 1 | 2.35e-1 |
| p.T120I | 1 | 5.84e-3 |
| p.R147S | 2.01e-4 | 1 |
| p.R183W | 1 | 8.08e-3 |
| p.R196C | 1 | 1.18e-1 |
| p.R196H | 1 | 3.43e-1 |
| p.T203M | 1.76e-7 | 1 |

| Related to Figure 3B, Kruskall-Wallis and Dunn test with adjusted p-value and no extreme outliers. | | |
|---|---|---|
| | WT | *rey021* |
| *rey021* | 3.60e-6 | – |
| p.L65V | 1.15e-2 | 1 |
| p.G74S | 1 | 3.11e-1 |
| p.T120I | 1 | 5.99e-3 |
| p.R147S | 7.34e-5 | 1 |
| p.R183W | 1 | 2.93e-3 |
| p.R196C | 1 | 2.50e-1 |
| p.R196H | 1 | 4.69e-1 |
| p.T203M | 4.25e-8 | 1 |

| Related to Figure 3D, ANOVA and Tukey_hsd with adjusted p-value and extreme outliers. | | | | | | |
|---|---|---|---|---|---|---|
| | p.L65V | p.G74S | p.T120I | p.R196C | p.R196H | p.T203M |
| WT | 3.03e-2 | 2.02e-1 | 1 | 9.91e-1 | 8.54e-1 | 3.07e-1 |

| Related to Figure 3D, Kruskall-Wallis and Dunn test with adjusted p-value and no extreme outliers. | | | | | | |
|---|---|---|---|---|---|---|
| | p.L65V | p.G74S | p.T120I | p.R196C | p.R196H | p.T203M |
| WT | 2.60e-1 | 3.93e-1 | 1 | 1 | 1 | 1 |

| Related to Figure 5A Kruskall-Wallis and Dunn test with adjusted p-value and extreme outliers. | | |
|---|---|---|
| | WT | *unc-2*(*gk366*) |
| *unc-2*(*gk366*) | 6.67e-3 | – |
| *rey021* | 5.69e-2 | 1 |
| p.L65V | 1 | 7.67e-3 |
| p.G74S | 1 | 1 |
| p.T120I | 4.25e-2 | 1 |
| p.R147S | 1 | 1 |
| p.R183W | 9.14e-1 | 1 |
| p.R196C | 1.66e-1 | 1 |
| p.R196H | 1.52e-3 | 1 |
| p.T203M | 9.15e-1 | 1 |

| Related to Figure 5A Kruskall-Wallis and Dunn test with adjusted p-value and no extreme outliers. | | |
|---|---|---|
| | WT | *unc-2*(*gk366*) |
| *unc-2*(*gk366*) | 6.16e-3 | – |
| *rey021* | 5.62e-2 | 1 |
| p.L65V | 1 | 7.02e-3 |
| p.G74S | 1 | 8.61e-1 |
| p.T120I | 4.05e-2 | 1 |
| p.R147S | 1 | 1 |
| p.R183W | 9.5e-1 | 1 |
| p.R196C | 1.56e-1 | 1 |
| p.R196H | 1.26e-3 | 1 |
| p.T203M | 8.75e-1 | 1 |
