## Supplement Movies for "Multiscale characterization of *Caenorhabditis elegans* mutants to probe functional mechanisms of human actin pathological variants": Legend.docx

Movie 1 :

Cytokinesis arrest, p.R196C embryo. Scale bar = 10µm.

Movie 2:

Gastrulation arrest, p.G74S embryo. Scale bar = 10µm.

Movie 3:

Ventral enclosure arrest, *rey021* embryo. Scale bar = 10µm.

Movie 4:

Extremity burst arrest, p.L65V embryo. Scale bar = 10µm.

Movie 5:

Paralyzed arrest, *rey021* embryo. Scale bar = 10µm.

Movie 6:

Cytokinesis defects, p.R196C embryo. LifeAct::mKate2 (Magenta), NMY-2::GFP (Green). Scale bar = 10µm. Anterior on the left.

Movie 7:

Meiosis defects, p.R196C embryo. LifeAct::mKate2 (Magenta), NMY-2::GFP (Green). Scale bar = 10µm. Anterior on the left.

Movie 8:

Bleb defects, p.T120I embryo. LifeAct::mKate2 (Magenta), NMY-2::GFP (Green). Scale bar = 10µm. Anterior on the left.
